# Molecular flexibility of DNA as a key determinant of RAD51 recruitment

**DOI:** 10.1101/392795

**Authors:** Federico Paoletti, Afaf El-Sagheer, Jun Allard, Tom Brown, Omer Dushek, Fumiko Esashi

## Abstract

The timely activation of homologous recombination is essential for the maintenance of genome stability, in which the RAD51 recombinase plays a central role. Biochemically, human RAD51 polymerises faster on single-stranded DNA (ssDNA) compared to double-stranded DNA (dsDNA), raising a key conceptual question: how does it discriminate between them? In this study, we tackled this problem by systematically assessing RAD51 binding kinetics on ssDNA and dsDNA differing in length and flexibility using surface plasmon resonance. By fitting detailed polymerisation models informed by our experimental datasets, we show that RAD51 is a mechano-sensor that exhibits a larger polymerisation rate constant on flexible ssDNA compared to rigid ssDNA or dsDNA. This model presents a new general framework suggesting that the flexibility of DNA, which may increase locally as a result of DNA damage, plays an important role in rapidly recruiting repair factors that multimerise at sites of DNA damage.

## Introduction

DNA double-strand breaks (DSBs) are cytotoxic lesions that can lead to chromosomal breaks, genomic instability and tumourigenesis in mammalian cells (Tubbs and Nussenzweig, 2017). Homologous recombination (HR) can offer an error-free DNA repair mechanism to restore genetic information at DSB sites, and in this way, contribute to genome stability. During HR-mediated repair, single-stranded DNA (ssDNA) overhangs are generated and rapidly coated with the ssDNA-binding replication protein A (RPA) (Chen and Wold, 2014). ssDNA-bound RPA is then exchanged for RAD51, the central ATP-dependent recombinase that catalyses HR-mediated repair. RAD51 polymerises on ssDNA to form a ssDNA-nucleoprotein filament and guide homologous strand invasion and DSB repair (Baumann et al., 1996).

The central mechanism of HR is evolutionarily highly conserved, and the bacterial RAD51 ortholog RecA has clear preference to polymerise on ssDNA over dsDNA (Benson et al., 1994). However, human RAD51 shows weaker binding affinity to ssDNA compared to RecA, and the mechanism by which RAD51 polymerises on ssDNA in preference to dsDNA remains enigmatic. Earlier studies using electrophoretic mobility shift assay (EMSA) have suggested that RAD51 binds both ssDNA and dsDNA with similar affinities (Benson et al., 1994), albeit its preferential ssDNA binding was visible in the presence of ammonium sulphate (Shim et al., 2006). These endpoint assays, however, do not provide information as to whether binding kinetics may contribute to a potential RAD51-dependent ssDNA / dsDNA discrimination mechanism. Indeed, more recent kinetic studies have revealed that RAD51 polymerisation consists of two phases: a rate-limiting nucleation phase and a growth phase (Hilario et al., 2009; Miné et al., 2007; Van der Heijden et al., 2007). A minimal polymer nucleus, with a length of either two-to-three or four-to-five RAD51 protomers (Hilario et al., 2009; Van der Heijden et al., 2007), is proposed to elicit the growth phase of RAD51 polymerisation. Intriguingly, RAD51 was shown to display faster association kinetics on ssDNA (Candelli et al., 2014) and slower dissociation kinetics on dsDNA, indicating that the RAD51-dsDNA complex is stable once formed (Miné et al., 2007). However, the mechanism underlying these kinetic differences is unclear.

A key difference between ssDNA and dsDNA is their molecular flexibility: ssDNA is known to be more flexible compared to dsDNA. The flexibility of a DNA molecule can be characterised by its persistence length (L_p_), a mechanical parameter quantifying polymer rigidity: the higher the persistence length, the more rigid the polymer. In the presence of monovalent or divalent salt, dsDNA displays an L_p_ of ∼ 30 - 55 nm (Baumann et al., 1997; Brunet et al., 2015), while ssDNA is much more flexible, with an L_p_ of 1.5 - 3 nm (Chi et al., 2013; Kang et al., 2014; Murphy et al., 2004). These observations imply that ssDNA can explore a much larger configurational space compared to dsDNA. It follows that the formation of a structured (less flexible) RAD51 polymer on ssDNA will incur a larger entropic energy penalty compared to the formation of a RAD51 polymer on dsDNA. Despite the clear thermodynamic implications of RAD51 polymerisation on DNA, the impact of DNA flexibility on RAD51 nucleoprotein filament formation has been largely overlooked.

In this study, we describe how RAD51 polymerises on DNA. Using a combination of surface plasmon resonance (SPR) and small-angle x-ray scattering (SAXS), we have assessed the RAD51 binding kinetics on DNA using ssDNA and dsDNA oligonucleotides differing in length and flexibility. Analyses of the SPR data using biochemical mathematical models revealed that RAD51 polymerisation required a minimal nucleus of four and two molecules on ssDNA and dsDNA, respectively. Interestingly, our analyses further uncovered that RAD51 is a mechano-sensor that polymerises faster on flexible DNA. We hypothesised that the entropic penalty to polymerisation on flexible DNA is offset by a large enthalpic contribution and show this to be the case by perturbing the RAD51 protomer-protomer interaction. We propose the DUET model (**<u>D</u>**na molec**<u>U</u>**lar fl**<u>E</u>**xibili**<u>T</u>**y), which explains the ability of RAD51 to preferentially and stably polymerise on highly flexible ssDNA compared to stiff ssDNA and dsDNA, enabled by the enthalpic energy of RAD51 polymerisation.

## Results

### RAD51 preferential binding to ssDNA is dependent on the length of DNA template

To evaluate how human RAD51 discriminates between ssDNA and dsDNA, we used SPR to assess the binding kinetics of untagged recombinant human wild-type (WT) RAD51 to a 50-mer mixed base ssDNA molecule (dN-50) and a 50-mer mixed base paired dsDNA molecule (dN-50p). These analyses performed in the presence of ATP and Ca^2+^, which blocks RAD51 ATP hydrolysis (Bugreev and Mazin, 2004), showed 1) faster RAD51 association to ssDNA compared to dsDNA, and 2) similar RAD51 lifetimes on both ssDNA and dsDNA (Fig. S1 A **and** B). These observations confirm previous findings that WT RAD51 distinguishes ssDNA from dsDNA through faster polymerisation on ssDNA (Candelli et al., 2014), validating SPR as a sensitive experimental system to determine RAD51 binding kinetics to DNA.

We then aimed to define the initial steps of RAD51 association with ssDNA. To this end, we generated a series of mixed base short ssDNA oligos with varying lengths, each of which can be bound by a restricted number of RAD51 molecules. As a single RAD51 molecule engages with three nucleotides of DNA (Short et al., 2016), a ssDNA consisting of 5 nucleotides (dN-5), 8 nucleotides (dN-8), 11 nucleotides (dN-11), 14 nucleotides (dN-14) and 17 nucleotides (dN-17) can accommodate up to one, two, three, four and five RAD51 molecules, respectively (Fig. 1A). Our systematic SPR measurements of WT RAD51 binding kinetics revealed no RAD51 binding to the dN-5, dN-8 and dN-11, slow association and moderate dissociation with dN-14, and slightly faster association and slow dissociation with dN-17 (Fig. 1 A **and** B). These observations suggested that three or less RAD51 molecules are unable to generate a stable nucleus on ssDNA, four RAD51 molecules form a quasi-stable nucleus, and five or more RAD51 molecules are able to form a highly stable nucleus.

**Figure 1.**
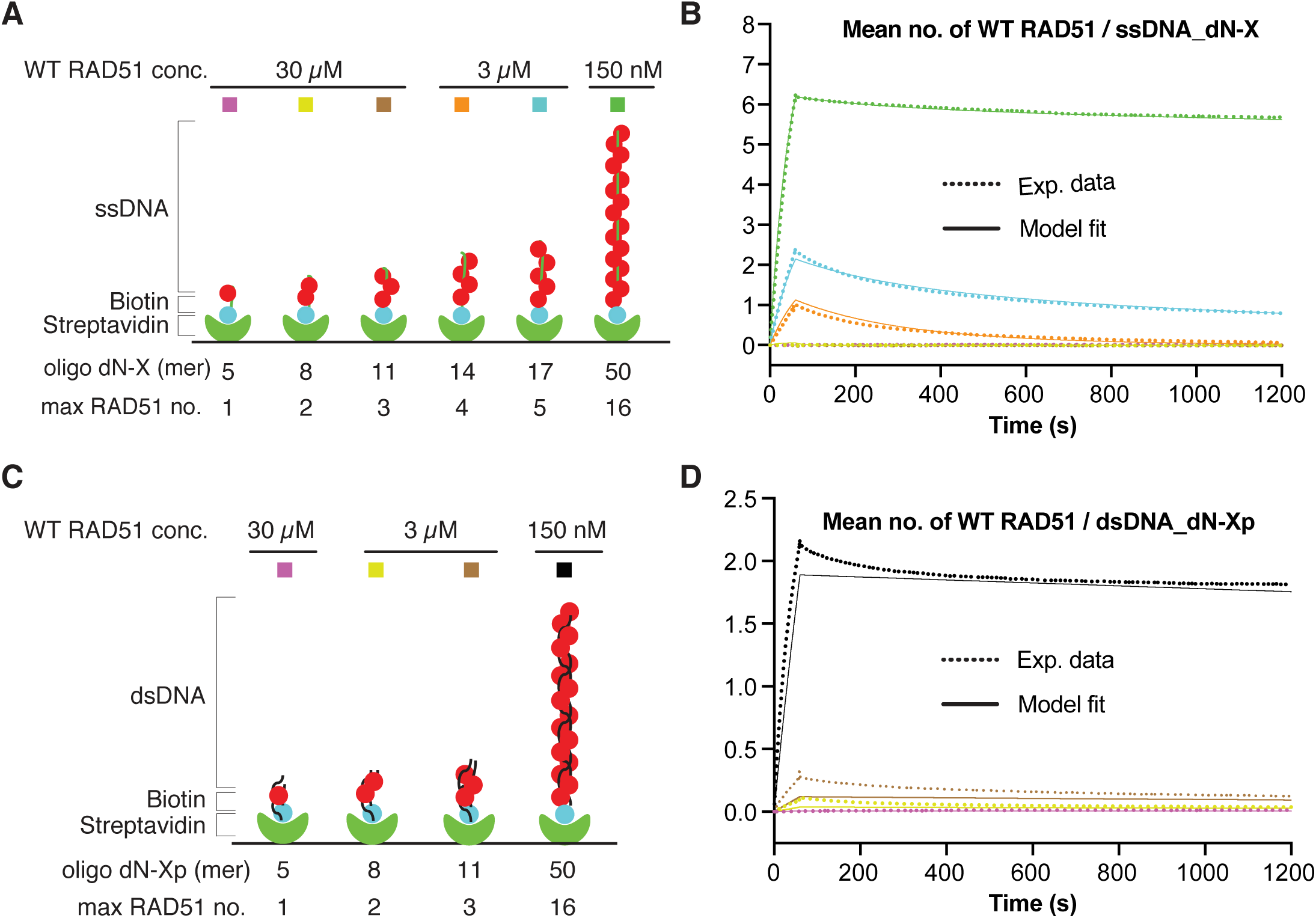
Kinetics of WT RAD51 binding to ssDNA and dsDNA oligos of varying length. ***A***. Biotinylated ssDNA oligos of indicated lengths were separately immobilised onto SPR CM5 chips *via* biotin-streptavidin interaction. Wild-type (WT) RAD51 was injected at the indicated concentrations to measure association and dissociation kinetics. ***B***. The dotted and solid curves respectively show the binding of RAD51 to the ssDNA oligos measured by SPR and the ODE model fits (see Fig. 2A for model description). ***C***. As in *A*, except biotinylated dsDNA molecules of indicated lengths were used to measure association and dissociation kinetics. ***D***. The dotted and solid curves respectively show the binding of RAD51 to the dsDNA oligos measured by SPR and the ODE model fits (see Fig. 2B for model description).

To similarly evaluate the initial steps of RAD51 association with dsDNA, we generated an analogous series of mixed base dsDNA consists of 5 base pairs (dN-5p), 8 base pairs (dN-8p) and 11 base pairs (dN-11p), which can accommodate up to one, two or three RAD51 molecules, respectively (Fig. 1C). Our SPR measurement detected no RAD51 binding to the dsDNA dN-5p, as was the case for the ssDNA dN-5. To our surprise, however, we detected slow association and moderate dissociation of WT RAD51 with the dN-8p, and even faster association and slower dissociation with the dN-11p (Fig. 1D). These observations initially suggested that one RAD51 molecule is unable to associates stably with dsDNA, but two RAD51 molecules form a quasi-stable nucleus, and three or more RAD51 molecules are able to form a highly stable nucleus. This suggested that WT RAD51 nucleation on dsDNA requires only 2-3 RAD51 molecules versus 4-5 on ssDNA. Altogether, these observations supports the idea that WT RAD51 can bind more strongly to short (≤ 11 bp) dsDNA compared to short (≤ 11 nt) ssDNA molecules, but binds more strongly to long (50 nt) ssDNA molecules compared to long (50 bp) dsDNA molecules.

### RAD51 polymerisation on ssDNA is promoted by faster adsorption and / or elongation compared to dsDNA

Informed by these SPR datasets, we developed ODE models describing RAD51 polymerisation on ssDNA and dsDNA, that could be simultaneously fit to all corresponding SPR curves. This approach would enable us to determine the mechanistic differences in RAD51 polymerisation kinetics on ssDNA and dsDNA.

The polymerisation of RAD51 in solution was based on mass action kinetics with a maximum length of 16 (see Supplementary Information for details). The model allows any RAD51 n-mer (1 ≤ n < 16) to associate with any other RAD51 m-mer (1 ≤ m < 16) to form an (m + n)-mer (1 < m + n ≤ 16). In addition, any RAD51 k-mer can fall apart in every possible combination of m-mers and n-mers (k = n + m) (e.g. a pentamer can fall apart to form a monomer and a tetramer, or a dimer and a trimer). This model provides the concentrations of RAD51 polymers in solution, and is solved in the steady-state so that it only depends on a single fit parameter K_D_ (i.e. the RAD51 promoter-promoter dissociation constant). The predicted concentrations are then inserted into the kinetic models to describe RAD51 polymerisation on ssDNA and dsDNA.

The next two kinetic models described the sequential formation of RAD51 polymers on ssDNA and on dsDNA. We tested models of increasing complexity based on mass action kinetics until we identified models that were able to fit the experimental datasets (see Supplementary Information for details). Both the ssDNA and dsDNA models allow for the adsorption of any RAD51 n-mer from solution onto DNA (provided at least n DNA binding sites are available), given that n-mers (1 ≤ n ≤ 16) can exist in solution. Polymer elongation occurs when a RAD51 m-mer in solution binds to a DNA-bound n-mer to form a DNA-bound (n+m)-mer, provided that at least n + m DNA binding sites are available and 1 < m + n ≤ 16. However, unbinding was assumed to only take place via single protomer dissociation and via the dissociation of short RAD51 nuclei. This is a valid assumption when considering that in all the SPR experiments in this study ATP hydrolysis is inhibited via the addition of Ca^2+^, leading to slow RAD51 filament disassembly which can be approximated as disassembly via monomer removal or via the removal of short, unstable RAD51 nuclei. The ssDNA model consists of a polymerisation forward rate constant (k_p_), an unstable reverse rate constant (k_u_), a quasi-stable reverse rate constant (k_q_), and a stable reverse rate constant (k_s_). k_p_ describes the adsorption and elongation of RAD51 polymers on ssDNA up to a maximum length of 16, k_u_ describes the dissociation of unstable nuclei (1-3 RAD51 molecules), k_q_ describes the dissociation of quasi-stable nuclei (4 RAD51 molecules), and k_s_ describes the dissociation of single RAD51 protomers (i.e. RAD51 monomers of a RAD51 polymer) (Figs. 1A **and** 2A). Similarly, kinetic model for dsDNA includes k_p_, k_u_, and k_s_, but without the need for k_q_ to describe dissociation of quasi-stable nuclei (i.e. 2 RAD51 molecules) (Fig. 1C). For dsDNA, k_u_ describes the rate of dissociation of RAD51 monomers not bound to a RAD51 polymer (Fig. 2B).

**Figure 2.**
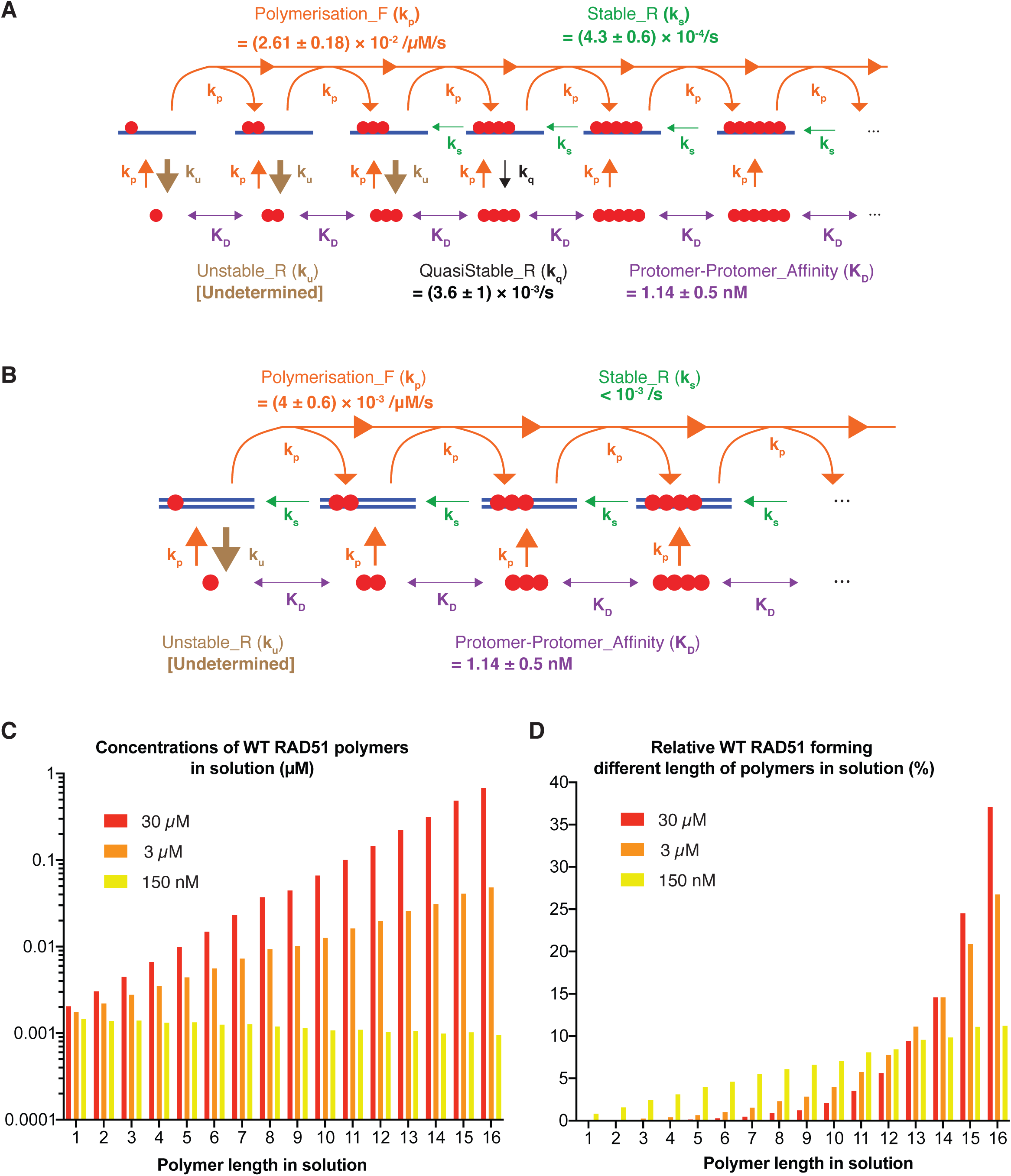
Mathematical models describing WT RAD51 polymerisation kinetics on ssDNA and dsDNA. ***A*.** Kinetic representation of the ordinary differential equation (ODE) model describing WT RAD51 polymerisation on ssDNA, consisting of five parameters: k_p_ (polymerisation forward rate), k_u_ (unstable reverse rate), k_q_ (quasi-stable reverse rate), k_s_ (stable reverse rate) and K_D_ (protomer-protomer interaction affinity). K_D_ predicts the concentrations of RAD51 polymers of variable length in solution, while k_p,_ k_u,_ k_q and_ k_s_ predict the speed of formation of RAD51 polymers on ssDNA. ***B***. Kinetic representation of the ODE model describing WT RAD51 polymerisation on dsDNA, consisting of four parameters: k_p_, k_u_, k_s_, K_D_. K_D_ predicts the concentrations of RAD51 polymers of variable length in solution, and k_p,_ k_u and_ k_s_ predict the speed of formation of RAD51 polymers on dsDNA. In panels *A* and *B*, purple arrows depict a simplified cartoon representation of RAD51 polymerisation in solution. The model allows for any RAD51 n-mer (1 ≤ n < 16) to associate with any other RAD51 m-mer (1 ≤ m < 16) to form an (m+n)-mer (1 < m + n ≤ 16), and any RAD51 k-mer to dissociate into any combination of n-mers and m-mers (k = n + m). The ssDNA and dsDNA models were calibrated by simultaneously fitting the SPR curves of ssDNA dN-X (dN-8, dN-14, dN-17, dN-50) and dsDNA dN-Xp (dN-5p, dN-8p, dN-11p, dN-50p) using the mode ABC-SMC particles. Mean values ± 1 SD of 3 mode ABC-SMC particles derived from model fits of 3 dN-X ^and dN-Xp repeats (n = 3). ssDNA k^_p_, ^k^_u_, ^k^_q_ and ^k^_s_ were fit to the ssDNA SPR curves, and dsDNA k_p,_ k_u and_ k_s were fit to the dsDNA SPR curves._ A single K_D_ was fit to all curves. ***C***. Predicted concentrations of WT RAD51 polymers in solution at equilibrium as a function of WT RAD51 monomer concentration (150 nM, 3 µM, 30 µM), prior to RAD51 injection onto DNA-coated SPR CM5 chips. ***D***. Predicted % WT RAD51 within each polymer state in solution at equilibrium as a function of WT RAD51 monomer concentration (150 nM, 3 µM, 30 µM), prior to RAD51 injection onto DNA-coated SPR CM5 chips. In panels *C* and *D*, the K_D_ was fixed to the mean value identified in Figs. 2 A **and** B (i.e. K_D_ = 1.14 nM), and the concentration of WT RAD51 monomer concentration was varied accordingly. In panel *D*, the % WT RAD51 values were calculated by multiplying the polymer concentrations by their respective polymer length, and dividing each value by the total RAD51 monomer concentration (i.e. % WT RAD51 = [*n*-mer] * *n* / [WT RAD51_monomer_]). [WT RAD51_monomer_] was calculated via coomassie staining image quantification using a BSA standard curve. It is important to note that the assumed maximum polymer length in solution (16-mer) leads to an overestimation of the concentration of each n-mer in solution, given that RAD51 is likely to form polymers of length greater than 16 at 150 nM, 3 µM and 30 µM WT RAD51 monomer concentration. However, this overestimation is likely to not alter the conclusion that WT RAD51 forms long polymers in solution at 150 nM - 30 µM concentration.

Using ABC-SMC to fit these models to the ssDNA dN-X data (Fig. 1B) and the dsDNA dN-Xp data (Fig. 1D), values for the model parameters were determined (Fig. 2 A **and** B, S2). This analysis identified a nano-molar range RAD51 protomer-protomer dissociation constant (dN-X and dN-Xp K_D_ = 1.14 ± 0.5 nM) as a key factor driving rapid RAD51 polymerisation on both ssDNA and dsDNA, similarly to other studies using either pressure perturbation fluorescence spectroscopy (Schay et al., 2016) or single molecule fluorescence microscopy (Candelli et al., 2014). This is due to the fact that this K_D_ value enables WT RAD51 to form abundant long polymers in solution at both 150 nM and 3 µM [WT RAD51] (Fig. 2 C **and** D). These RAD51 polymers can then directly adsorb onto DNA and elongate, therefore enabling WT RAD51 to polymerise faster on the dN-50 and dN-50p (150 nM [WT RAD51]) compared to the dN-14, dN-17, dN-11p and dN-8p oligos (3 µM [WT RAD51]) despite a 20-fold lower RAD51 injection concentration.

Importantly, we also identified a 6.5-fold higher ssDNA polymerisation forward rate constant (dN-X k_p_ = (2.6 ± 1.8) × 10^-2^ /µM/s) compared to dsDNA (dN-Xp k_p_ = (4 ± 0.6) × 10^-3^ /µM/s), explaining the overall faster RAD51 polymerisation on ssDNA (Fig. 2 A **and** B). Overall, these analyses suggest that WT RAD51 adsorption and/or elongation is faster on ssDNA compared to dsDNA, and that a high protomer-protomer affinity enables WT RAD51 to nucleate and elongate effectively on ssDNA and dsDNA even at low concentrations of RAD51.

### RAD51 polymerises faster on flexible DNA

Overall, our analyses suggest that WT RAD51 can nucleate more efficiently on short dsDNA molecules but polymerises significantly faster on long ssDNA molecules. We speculate that the difference in RAD51 polymerisation speed is due to the higher flexibility of ssDNA compared to dsDNA. Explicitly, we propose a Bend-To-Capture (BTC) mechanism to explain how DNA flexibility can impact polymerisation kinetics (Fig. 3A). A free RAD51 monomer or polymer in solution would need to generate two sequential, non-covalent interactions to be incorporated into the growing polymer: the interaction with the exposed interface of an existing RAD51 polymer and with the exposed scaffold DNA. To avoid steric clashes, we reasoned that each of these interactions requires naked DNA immediately next to the existing RAD51 polymer in a configuration that can bend away from the preferred direction of the polymer. In this way, flexible DNA is expected to explore more conformations compatible with the further addition of RAD51 per unit time.

**Figure 3.**
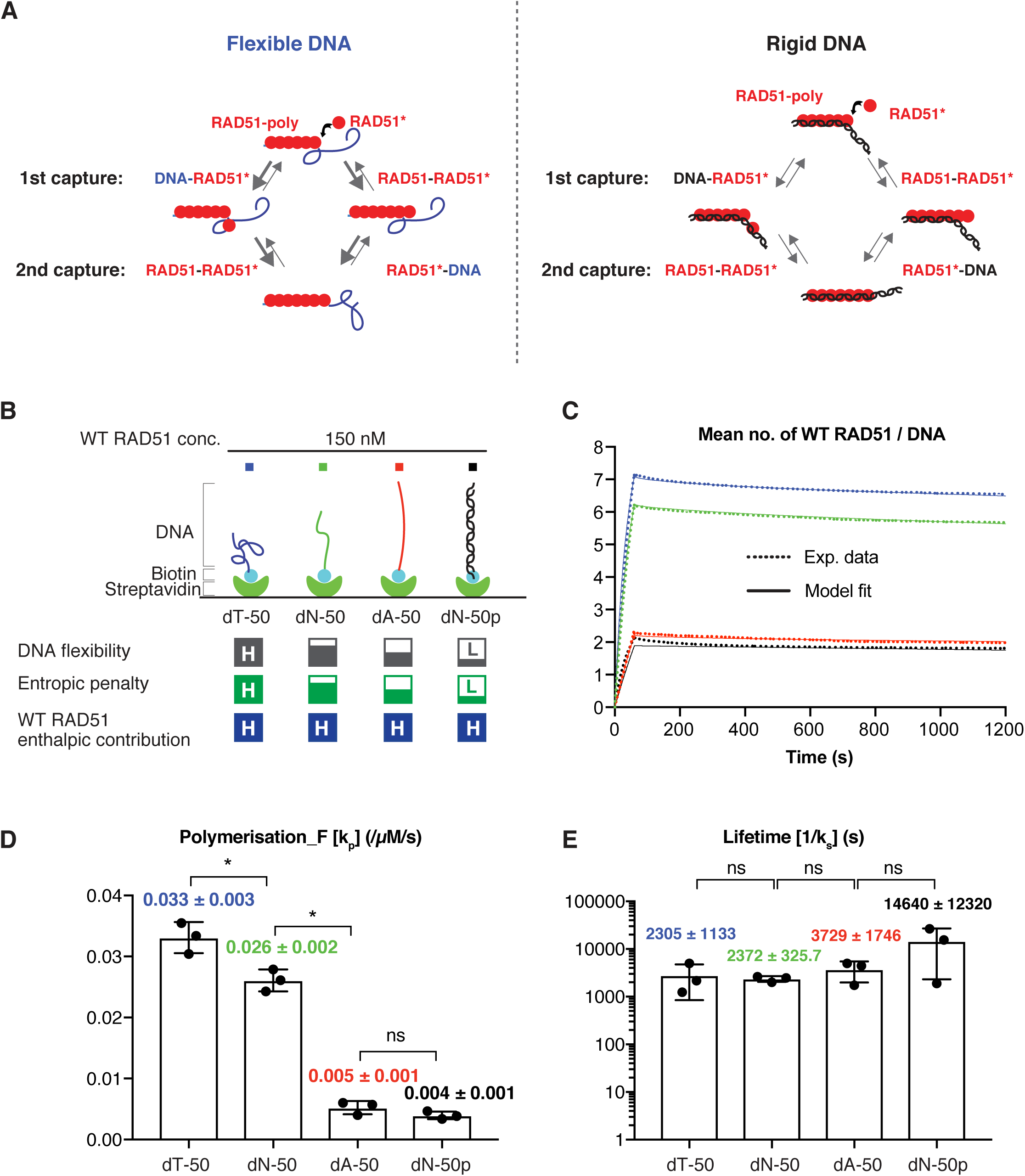
Kinetics of WT RAD51 binding to DNA of varying flexibility. ***A***. A depiction of the Bend-to-Capture (BTC) mechanism. RAD51 generates two sequential, non-covalent interactions (a RAD51 protomer-protomer interaction and a RAD51-DNA interaction, or *vice versa*) faster on flexible DNA (depicted as ssDNA) compared to rigid DNA (depicted as dsDNA). The free RAD51 monomer or polymer in solution (here depicted as a monomer) to be incorporated into the growing polymer is shown with an asterisk. ***B***. Biotinylated DNA molecules of varying flexibility were separately immobilised onto SPR CM5 chips *via* biotin-streptavidin interaction, and WT RAD51 was injected at the indicated concentrations to measure association and dissociation kinetics. The flexibility of respective DNA, the expected entropic penalties upon RAD51 binding to corresponding DNA and enthalpic contribution of WT RAD51 are indicated in gray, green and blue boxes. H and L in each box denote high and low, respectively. ***C***. WT RAD51 SPR curves for respective DNA oligos (dotted lines) and ODE model fits (solid lines). ***D* and *E***. Bar plots of fitted k_p_ values (*D*) and k_s_ values (*E*) for each SPR curves. In panel *D*, all parameters except k_p_ were fixed to the mean values as identified in Fig. 2 A **and** B, and k_p_ was fitted using lsqcurvefit (MATLAB). In panel *E*, all parameters except k_s_ and k_p_ were fixed to the mean values as identified in Fig. 2 A **and** B, and k_s_ and k_p_ were fitted using lsqcurvefit (MATLAB). Only the k_s_ values are reported here. Mean ± 1 SD of 3 ODE model fits (n=3). *D*: un-paired, one-tailed Mann-Whitney-Wilcoxon tests. *E*: un-paired, two-tailed Mann-Whitney-Wilcoxon tests. * p ≤ 0.05; ns = non-significant.

To test this notion, we designed an experiment to measure the kinetics of WT RAD51 binding to DNA oligos of varying flexibility. It has been shown that poly-dT ssDNA, which is widely used for RAD51 binding assays, is highly flexible, while poly-dA ssDNA is highly rigid due to base stacking interactions (Mills et al., 1999) (Sim et al., 2012). Consistently, our SAXS-derived persistence length measurements showed ssDNA dT-50 is the most flexible oligo, followed by dN-50, dA-50 and dsDNA dN-50p (Fig. S3 **and** Table 1). By measuring RAD51 binding kinetics to these DNA oligos (Fig. 3B), we found that WT RAD51 indeed displayed faster association to the dT-50 compared to the dN-50, and the model fit suggests this is due to a higher polymerisation rate constant (k_p_) (Fig. 3 C **and** D). Furthermore, WT RAD51 displayed very slow association to the dA-50, comparable to that of dsDNA dN-50p, which is explained by a lower polymerisation rate constant (Fig. 3 C **and** D). Conversely, individually fitting the unstable reverse rate constant (k_u_), the quasi-stable reverse rate constant (k_q_) or the stable reverse rate constant (k_s_) cannot explain the ssDNA dT-50 or dA-50 data (Fig. S4 A-C). In addition, for RAD51 DNA-bound polymers of length greater than 3, the fractions of RAD51 in each polymer state are similar on ssDNA dT-50, dN-50, dA-50 and dsDNA dN-50p, suggesting WT RAD51 forms a similar proportion of polymers length 4 - 16 across the four DNA molecules (Fig. S4D). Finally, we observed that the lifetime of DNA-bound RAD51 protomers was independent of the flexibility of the underlying DNA (Fig. 3E). Taken together, these observations suggest that although WT RAD51 is able to form a stable nucleus with fewer molecules on dsDNA compared to ssDNA (two versus five), it elongates and/or adsorbs more efficiently on ssDNA because of its higher flexibility, consistent with our proposed BTC mechanism.

**Table 1.** Persistence lengths (L_p_) ± 95% confidence intervals are estimated from fitted Kuhn lengths for the ssDNA dN-50, dA-50, dT-50 and the dsDNA dN-50p SAXS plots. All data manipulation and model fitting were done using SasView. It is likely that the relatively low dsDNA dN-50p L_p_ of 7.34 nm compared to the average dsDNAp L_p_ of 30 - 55 nm (Baumann et al., 1997; Brunet et al., 2015) is due to the low GC content of the dN-50p (36%).

| Oligo Name | Kuhn length (nm) | Persistence length (nm) | Persistence length error (95% CI) (nm) |
| --- | --- | --- | --- |
| ssDNA dT-50 | 2.6702 | 1.3351 | 0.200185 |
| ssDNA dN-50 | 4.5177 | 2.25885 | 0.048095 |
| ssDNA dA-50 | 9.3488 | 4.6744 | 0.13126 |
| dsDNA dN-50p | 14.674 | 7.337 | 0.07379 |

**Table 2.**
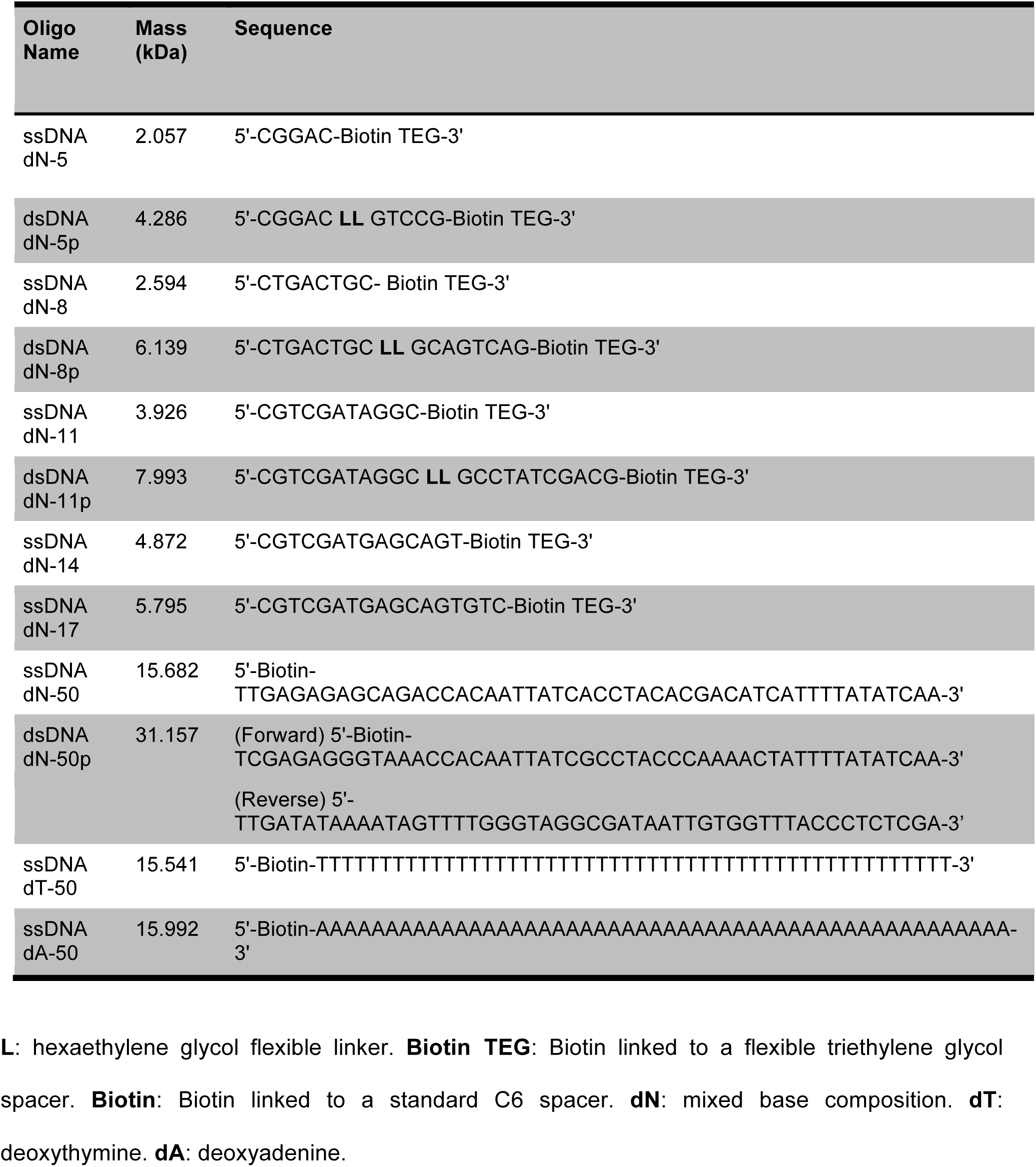
The list of DNA oligos used in this study.

### RAD51 protomer-protomer interaction compensates for the entropic penalty of flexible DNA

So far, we have observed that WT RAD51 associates faster to flexible ssDNA compared to dsDNA, and the proposed BTC mechanism describes how higher DNA flexibility can induce faster RAD51 polymerisation. However, the formation of a stiff RAD51 polymer on highly flexible DNA is expected to be associated with a large entropic penalty because RAD51 polymerisation stiffens ssDNA and dsDNA to a similar extent (L_p_ ∼ 200 nm for both RAD51-ssDNA and RAD51-dsDNA complexes)(Miné et al., 2007), therefore dramatically reducing the number of spatial configurations that a flexible DNA polymer can explore. RAD51 polymerisation on flexible ssDNA is expected to incur a higher entropic penalty compared to its polymerisation on stiff ssDNA or dsDNA (Fig. 3 B, “Entropic penalty”). Since we do not observe this in the WT data, we hypothesized that the enthalpic contribution of WT RAD51 polymerisation on DNA is sufficiently large to dominate any entropic penalty to associate with ssDNA and dsDNA (Fig. 3 B, “WT RAD51 enthalpic contribution”). However, if this enthalpic contribution was to be reduced, our model predicts that at equilibrium there would be less RAD51 bound to flexible DNA compared to stiff DNA. We define this as the Entropic Penalty Compensation (EPC) mechanism, which proposes that the favourable enthalpic energy of RAD51 polymerisation on DNA is sufficient to compensate for any DNA-dependent entropic penalty. Specifically, we predict that, upon reduction of the enthalpic contribution, RAD51 will exhibit faster unbinding from flexible DNA compared to stiff DNA, as this is consistent with both reduced RAD51 binding at equilibrium and faster binding due to the Bent-to-Capture mechanism (Fig. 4A).

**Figure 4.**
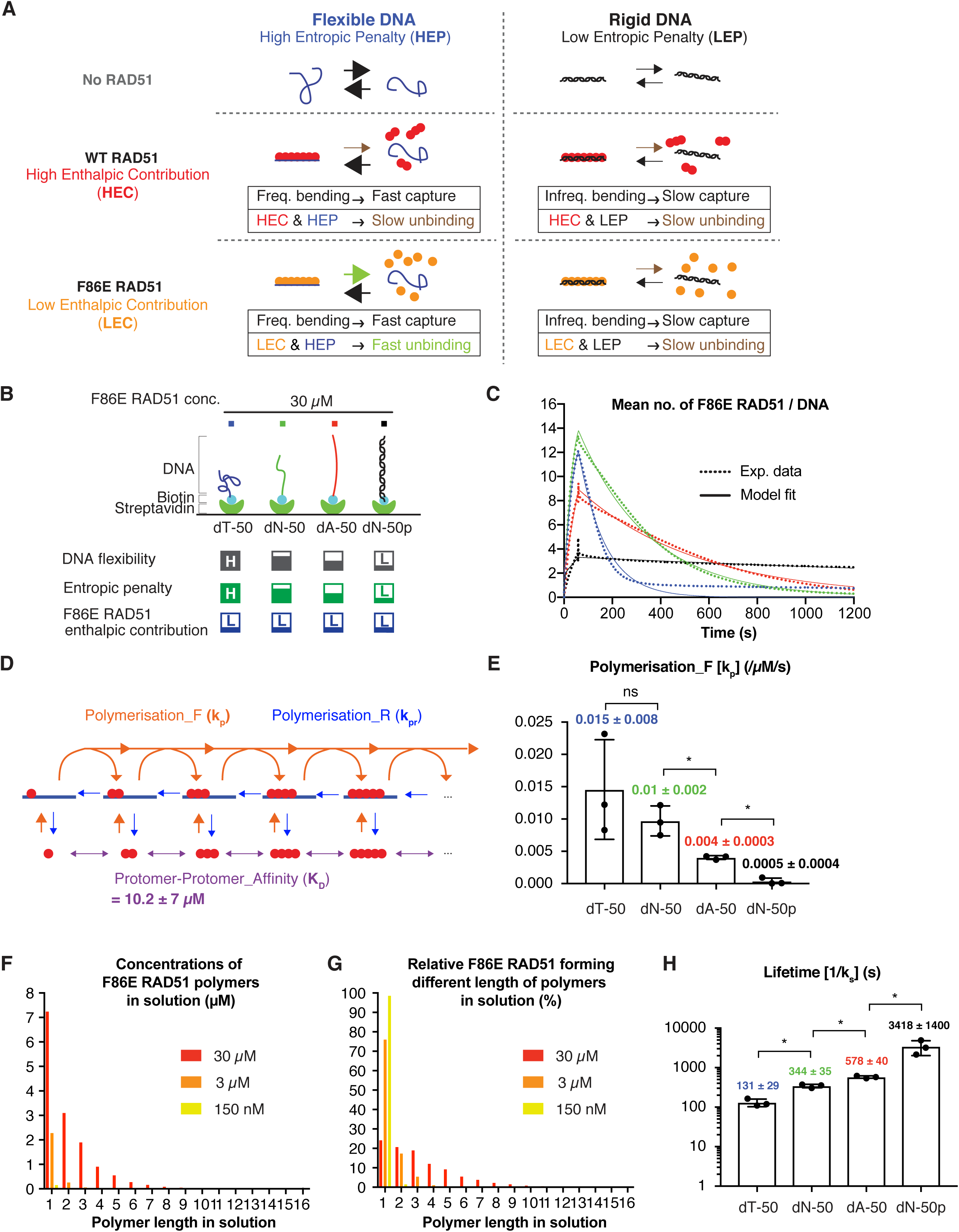
Kinetics of monomeric F86E RAD51 binding to DNA of varying flexibility. ***A***. A depiction of the Entropic Penalty Compensation (EPC) mechanism. The enthalpic contribution of WT RAD51 polymerisation enables RAD51 to overcome the entropic penalty to binding flexible DNA (here depicted as ssDNA). F86E RAD51 has a lower RAD51 protomer-protomer affinity and a consequent lower enthalpic contribution to polymerisation, and thus cannot overcome the entropic penalty for binding to flexible DNA. Both WT and F86E RAD51 can overcome the modest entropic penalty to binding stiff DNA (here depicted as dsDNA). Thick arrows indicate fast capture and fast unbinding, while thin arrows indicate slow capture and slow unbinding. ***B***. As in Fig. 3B, but F86E RAD51 was injected at the indicated concentration to measure association and dissociation kinetics. The flexibility of respective DNA, the expected relative entropic penalties upon F86E RAD51 binding to each DNA molecule and enthalpic contribution of F86 RAD51 are indicated in gray, green and blue boxes. H and L shown in each box indicate high and low, respectively. ***C*.** F86E RAD51 SPR curves (dotted lines) and corresponding model fits (solid lines) using the model shown in panel *D*. Using ABC-SMC, individual forward rates (k_p_) and reverse rates (k_pr_) were fitted to each experimental curve, while a single K_D_ was fitted to all experimental curves. ***D***. Kinetic representation of the ordinary differential equation (ODE) model describing F86E RAD51 polymerisation on ssDNA and dsDNA, consisting of three parameters: k_p_ (polymerisation forward rate), k_pr_ (polymerisation reverse rate), K_D_ (protomer-protomer dissociation constant). The K_D_ predicts the concentrations of F86E RAD51 polymers of variable length in solution, while k_p_ and k_pr_ predict the speed of formation of F86E RAD51 polymers on ssDNA and dsDNA. A simplified model compared to the models in Fig. 2 A **and** B was used for F86E RAD51 because the kinetics of F86E RAD51 to the dN-5, dN-5p, dN-8, dN-8p, dN-11, dN-11p, dN-14 and dN-17 were not systematically assessed due to low F86E purification yield. The predicted K_D_ value for F86E is shown. Mean value of mode particles ± 1 SD derived from the ABC-SMC fits (n=3). ***E***. Bar plots of fitted k_p_ values for the SPR curves. Mean values of mode particles ± 1 SD derived from the ABC-SMC fits (n=3). Un-paired, one-tailed Mann-Whitney-Wilcoxon tests. * p ≤ 0.05; ns = non-significant. ***F***. Predicted concentrations of F86E RAD51 polymers in solution at equilibrium as a function of F86E RAD51 monomer concentration (150 nM, 3 µM, 30 µM), prior to RAD51 injection onto DNA-coated SPR CM5 chips. ***G***. Predicted % F86E RAD51 within each polymer state in solution at equilibrium as a function of F86E RAD51 monomer concentration (150 nM, 3 µM, 30 µM), prior to RAD51 injection onto DNA-coated SPR CM5 chips. In panels *F* and *G*, the K_D_ was fixed to the mean value identified in Fig. 4D (i.e. K_D_ = 10.2 µM), and the concentration of F86E RAD51 monomer concentration was varied accordingly. In panel *G*, the % F86E RAD51 values were calculated by multiplying the polymer concentrations by their respective polymer length, and dividing each value by the total RAD51 monomer concentration (i.e. % F86E RAD51 = [*n*-mer] * *n* / [F86E RAD51_monomer_]). [F86E RAD51_monomer_] was calculated via coomassie staining image quantification using a BSA standard curve. ***H*.** Bar plots of fitted k_pr_ values for the SPR curves. Mean values of mode particles ± 1 SD derived from the ABC-SMC fits (n=3). Un-paired, one-tailed Mann-Whitney-Wilcoxon tests. * p ≤ 0.05; ns = non-significant.

To directly test this hypothesis, we assessed the kinetics of a RAD51 mutant, which confers a reduced enthalpic contribution compared to WT RAD51. We predicted that, with such a mutation, the stability of the mutant RAD51 polymers on DNA would become more dependent on the flexibility of the underlying DNA (Fig. 4 A **and** B). We took advantage of a phenylalanine to glutamate substitution at RAD51 residue 86 (F86E), which reduces the RAD51 protomer-protomer interaction affinity (Pellegrini et al., 2002; Esashi et al., 2007; Yu et al., 2003). Indeed, using size-exclusion chromatography with multi-angle light scattering (SEC-MALS), we confirmed that F86E RAD51 is primarily monomeric in solution (Fig. S5 A-C). We then used SPR to measure the binding kinetics of F86E RAD51 to ssDNA dT-50, dN-50, dA-50 and dsDNA dN-50p, and used ABC-SMC to fit a simplified polymerisation model simultaneously to all four data sets (Fig. 4D). Importantly, we fit a single reverse rate (k_pr_ = k_u_ = k_q_ = k_s_) to the F86E RAD51 SPR data, given that the F86E RAD51 binding kinetics to the short ssDNA oligos (dN-5, dN-8, dN-11, dN-14, dN-17) and dsDNA oligos (dN-5p, dN-8p, dN-11p) were not systematically measured due to low F86E purification yield. It was immediately evident that F86E RAD51 shows significantly reduced affinity for ssDNA and dsDNA compared to WT RAD51, with no detectable binding to DNA at the concentration of 150 nM or 3 µM (Fig. S5 D **and** E). Nonetheless, at 30 µM, we observed F86E RAD51 displays faster elongation on more flexible DNA (Fig. 4 B **and** E, Fig. S6), as we observed for WT RAD51 (Fig. 3 C **and** D). Additionally, DNA-binding at 30 µM can be explained by the fact that F86E RAD51 (K_D_ = 10.2 ± 7 µM) only forms abundant, nucleus-size polymers in solution (2-5 RAD51 molecules) at 30 µM [F86E RAD51] (Fig. 4 F **and** G). This is in contrast to WT RAD51 (K_D_ = 1.14 ± 0.5 nM), which can form abundant, long polymers in solution at 150 nM or 3 µM [WT RAD51] (Fig. 2 C **and** D). Strikingly, F86E RAD51 displayed reduced lifetimes (1/k_pr_) that inversely correlated with the flexibility of DNA (Fig. 4H). This observation is in sharp contrast with WT RAD51, which formed stable, long-lived polymers independently of DNA flexibility (Fig. 3E). Taken together, these observations support the notion that the enthalpic contribution of WT RAD51 polymer formation is sufficient to offset the large entropic penalty associated with polymerisation on flexible DNA. As a result, RAD51 is able to polymerise more rapidly on flexible DNA despite incurring a larger entropic penalty.

## Discussion

In this study, we have analysed the kinetics of RAD51 binding to ssDNA and dsDNA oligos of varying length and flexibility *via* a combination of SPR, SAXS and mathematical modelling to understand how RAD51 discriminates between ssDNA and dsDNA. Based on our observations, we propose the DUET (**<u>D</u>**na molec**<u>U</u>**lar fl**<u>E</u>**xibili**<u>T</u>**y) model, which describes how RAD51 polymerises on ssDNA in preference to dsDNA: 1) RAD51 has a faster polymerisation rate constant on flexible DNA because flexible DNA explores more conformations compatible with RAD51 binding per unit time (the BTC mechanism) (Fig. 3A), and 2) the enthalpic contribution of RAD51 polymerisation enables RAD51 to overcome the entropic penalty to binding to flexible DNA (the EPC mechanism) (Fig. 4A).

The diploid human genome consists of 6.4 billion base pairs and ∼ 50 endogenous DSBs are estimated to occur in every cell cycle (Vilenchik and Knudson, 2003). In human cells, the resection step of HR-mediated DSB repair can generate ssDNA overhangs up to 3.5 k nucleotides in length (Zhou et al., 2014), which then serve as a platform for RAD51 polymerisation. Given that ssDNA overhangs would constitute only ∼ 0.005 % of the total genomic DNA (3.5 k nucleotides x two overhangs x 50 DSBs / 6.4 billion base pairs), the polymerisation of RAD51 on resected ssDNA needs to be greatly directed. The average nuclear concentration of RAD51 is estimated at ∼ 100 nM (Reuter et al., 2014), and several RAD51 mediators, such as BRCA2, PALB2 and RAD52, contribute to increase the RAD51 local concentration at DSBs and/or RAD51 binding to ssDNA, while preventing its association with dsDNA (Buisson et al., 2010; Carreira and Kowalczykowski, 2011; Jensen et al., 2010; Ma et al., 2017a; Miyazaki et al., 2004; Zhao et al., 2015). Nonetheless, to date, *in vivo* evidence that these RAD51 mediators are sufficient to promote RAD51 polymerisation on DSB-derived ssDNA in preference to the bulk of undamaged dsDNA is limited.

It is important to note that resected ssDNA is first bound by RPA prior to RAD51 polymerisation. RPA is a heterotrimer complex with six-OB folds, four of which can associate tightly with ssDNA in a stepwise manner (Fan and Pavletich, 2012; Zou et al., 2006). RPA binding to ssDNA is believed to eliminate ssDNA secondary structures to facilitate RAD51 polymerisation. Significantly, previous kinetic studies have demonstrated a rapid exchange of ssDNA-bound RPA with free RPA available in solution, a phenomenon defined as ‘facilitated exchange’ (Ma et al., 2017b). Given that one RPA heterotrimer has a footprint of 30 nucleotides and one RAD51 protomer binds to 3 nucleotides of ssDNA (Short et al., 2016), the dissociation of a single RPA heterotrimer would expose enough ssDNA to accommodate up to ten RAD51 molecules. Hence, it is reasonable to speculate that RPA dissociation provides highly flexible ssDNA, which in turn can promote the adsorption and elongation steps of RAD51 polymerisation.

Another major unanswered question regarding RAD51 polymerisation is how RAD51 can distinguish ssDNA generated by DSB resection from ssDNA formed during normal cellular processes, such as DNA replication and transcription. Spontaneous RAD51 polymerisation on all ssDNA would be problematic, as it may trigger undesired, toxic recombination events or disruption of DNA replication and transcription, causing genomic instability. In this context, the DUET model is particularly appealing: ssDNA generated during transcription and replication does not have free ends and could therefore be less flexible compared to resected ssDNA with a free 3’ overhang. This in turn may limit RAD51 polymerisation on transcription- and replication-derived ssDNA, while promoting it on flexible ssDNA at DSBs. Intriguingly, a recent report demonstrated that poly(dA) stretches at replication forks are more vulnerable to DNA damage as they are unprotected by RPA, conforming early-replicating fragile sites (Tubbs et al., 2018). Our study further suggests that such poly(dA)-associated DNA damage is less efficiently repaired by RAD51-mediated HR, increasing genome instability at these loci.

It should be pointed out, however, that this study has built on the RAD51 kinetics detectable in the presence of Ca^2^+ which blocks RAD51-mediated ATP hydrolysis. This condition slows down RAD51 dissociation from DNA, hence enabling sensitive detection of RAD51 binding to short DNA substrates, which is otherwise challenging. Nonetheless, future studies to support our findings under conditions allowing RAD51’s ATP hydrolysis will shed further light on the dynamics of RAD51 polymerisation on DNA. Most significantly, this work paves the way for further work to include the impact of other HR mediator proteins, such as BRCA2 and RAD51 paralogs (Shahid et al., 2014; Taylor et al., 2016), to elaborate genome integrity control mechanisms.

Beyond HR repair, this study suggests a general framework to understand how DNA binding proteins are recruited to sites of DNA damage. Numerous DNA repair machineries are composed of multiple subunits with several binding interfaces to DNA and other co-factors. It is therefore tempting to speculate that broken DNA ends may enhance recruitment of such repair complexes simply due to increased DNA flexibility. In line with this notion, increased mobility of broken DNA has been demonstrated in mammalian cells (Aten et al., 2004; Cho et al., 2014; Aymard et al., 2017). Hence, this work presents a conceptual advancement in linking DNA repair and DNA flexibility, adding an important dimension which should be taken account of when assessing repair process both in biochemical assays and in cellular contexts.

## Material and methods

### RAD51 Mutagenesis

The bacterial expression vector pET11d (Merck-Millipore) carrying the human WT RAD51 (WT-pET11d) was used as a template for the PCR-mediated QuikChange site-directed mutagenesis (Agilent Technologies) to introduce F86E substitution (F86E-pET11d) with a forward primer 5’-GCTAAATTAGTTCCAATGGGTGAGACC ACTGCAACTGAATTCCACC - 3’ and a reverse primer 5’-GGTGGAATTCAGTTGCAGTGGTCT CACCCATTGGAACTAATTTAGC - 3’.

### RAD51 Protein Purification

To express RAD51, Rosetta 2 (DE3) pLysS cells (Novagen) carrying the WT-pET11d or F86E-pET11d were grown in LB media containing 100 µg/ml ampicillin and 25 µg/ml chloroamphenicol, and by adding 0.5 mM IPTG at OD_595_ = 0.6, protein expression was induced. Cell pellets were resuspended in PBS and mixed with the equal volume of lysis buffer (3 M NaCl, 100 mM Tris-HCl pH 7.5, 4 mM EDTA pH 8, 20 mM ß-mercaptoethanol, Sigma Protease Inhibitor Cocktail (Sigma)). The suspension was sonicated and spun at 20k rpm with a 45 Ti rotor (Beckman). The supernatant was slowly mixed with 0.1 % polyethylenimine at 4°C for 1 hour and spun at 20K rpm using a 45 Ti rotor to remove DNA. The supernatant was then slowly mixed with an equal volume of 4 M ammonium sulphate (2 M final concentration) at 4°C for 1 hour and spun at 10 K rpm using a JA-17 rotor (Beckman). Pellets containing RAD51 were suspended in 25 ml of resuspension buffer (0.5 M KCl, 50 mM Tris-HCl pH 7.5, 1 mM EDTA, 2 mM DTT, 10 % glycerol) and spun again at 20k rpm to remove residual DNA. The supernatant was then dialysed overnight at 4°C in TEG buffer (50 mM Tris pH 7.5, 1 mM EDTA, 2 mM DTT, 10 % glycerol) containing 200 mM KCl (TEG200). For F86E RAD51, the cell lysate was prepared as for WT RAD51, and dialysed overnight at 4°C in TEG buffer containing 50 mM KCl (TEG75).

RAD51 purification was carried out by chromatography at 4°C using the AKTA Pure Protein Purification System (GE Healthcare). For WT RAD51, the dialysed WT RAD51 containing sample was loaded onto a 5 ml HiTrap Heparin column (GE Healthcare) and eluted *via* a linear gradient of 200 mM – 600 mM KCl. The peak fractions were pooled and dialysed in TEG200, and concentrated on a 1 ml HiTrap Q column (GE Healthcare) followed by the isocratic elusion with TEG buffer containing 600 mM KCl (TEG600). Peak fractions containing WT RAD51 were dialysed in SPR buffer (150 mM KCl, 20 mM Hepes pH 7.5, 2 mM DTT, 10 % glycerol), aliquoted, snap frozen and stored at −80°C. Similarly, the dialysed F86E RAD51 containing cell lysate was fractionated through a 5 ml HiTrap Heparin column, but with a linear gradient of 75 mM – 600 mM KCl. The flow-through and F86E RAD51 peak fractions were pooled and dialysed in TEG buffer containing 50 mM KCl (TEG50) and reloaded onto a 5 ml HiTrap Heparin column. Following a 50 mM – 600 mM KCl linear gradient elution, F86E RAD51 peak fractions were pooled and dialysed in TEG buffer containing 100 mM KCl (TEG100). The sample was concentrated on a 1 ml HiTrap Q column followed by TEG600 isocratic elution. Peak fractions were applied on a 24 ml Superdex200 10/300 GL size-exclusion column (GE Healthcare), and fractions containing monometic F86E RAD51 were pooled and dialysed in TEG100. F86E RAD51 was then re-concentrated using a 1 ml HiTrap Q column and TEG600 isocratic elution. The peak fractions were dialysed in SPR buffer, aliquoted, snap frozen and stored at −80°C.

### Multi-Angle Light Scattering

To confirm the monomeric status of F86E RAD51, the peak size-exclusion chromatography F86E RAD51 elution fraction was serially diluted (5, 1:1 serial dilutions) and loaded onto a Superdex200 10/300 GL size exclusion column (GE Healthcare) equilibrated with MALS Buffer (150 mM KCl, 50 mM Tris pH 7.5, 1 mM EDTA, 2 mM DTT). Each elution was analysed using a Wyatt Heleos8+ 8-angle light scatterer linked to a Shimadzu HPLC system comprising LC-20AD pump, SIL-20A Autosampler and SPD20A UV/Vis detector. Data collection was carried out at the Department of Biochemistry, University of Oxford. Data analysis was carried out using the ASTRA software (Wyatt).

### DNA Oligo and Duplex Synthesis

The dN-5p, dN-8p and dN-11p dsDNA sequences were designed as two complimentary ssDNA sequences connected *via* two units of hexaethylene glycol (flexible linker). The dN-5, dN-8, dN-11 ssDNA molecules were designed using one of the two corresponding dsDNA annealing sequences. The dN-14 and dN-17 ssDNA molecules were designed by extending one of the two dN-11p dsDNA annealing sequences. Standard DNA phosphoramidites, solid supports, 3’-Biotin-TEG CPG and additional reagents were purchased from Link Technologies Ltd and Applied Biosystems Ltd. All oligonucleotides were synthesized on an Applied Biosystems 394 automated DNA / RNA synthesizer using a standard 1.0 *µ*mole phosphoramidite cycle of acid-catalyzed detritylation, coupling, capping, and iodine oxidation. Stepwise coupling efficiencies and overall yields were determined by the automated trityl cation conductivity monitoring facility and in all cases were >98.0%. All β-cyanoethyl phosphoramidite monomers were dissolved in anhydrous acetonitrile to a concentration of 0.1 M immediately prior to use. The coupling time for normal A, G, C, and T monomers was 60 s, and the coupling time for the hexaethylene glycol phosphoramidite monomer (from link) was extended to 600 s. Cleavage of the oligonucleotides from the solid support and deprotection was achieved by exposure to concentrated aqueous ammonia solution for 60 min at room temperature followed by heating in a sealed tube for 5 h at 55 °C. Purification of oligonucleotides was carried out by reversed-phase HPLC on a Gilson system using a Brownlee Aquapore column (C8, 8 mm x 250 mm, 300 Å pore) with a gradient of acetonitrile in triethylammonium bicarbonate (TEAB) increasing from 0% to 50% buffer B over 20 min with a flow rate of 4 ml/min (buffer A: 0.1 M triethylammonium bicarbonate, pH 7.0, buffer B: 0.1 M triethylammonium bicarbonate, pH 7.0 with 50% acetonitrile). Elution of oligonucleotides was monitored by ultraviolet absorption at 295 or 300 nm. After HPLC purification, oligonucleotides were freeze dried then dissolved in water without the need for desalting. All oligonucleotides were characterised by negative-mode HPLC-mass spectrometry using either a Bruker micrOTOFTM II focus ESI-TOF mass spectrometer with an Acquity UPLC system, equipped with a BEH C18 column (Waters) or a Waters Xevo G2-XS QT mass spectrometer with an Acquity UPLC system, equipped with an Acquity UPLC oligonucleotide BEH C18 column (particle size: 1.7 µm; pore size: 130 Å; column dimensions: 2.1 x 50 mm). Data were analysed using Waters MassLynx software or Waters UNIFI Scientific Information System software.

### Small Angle X-Ray Scattering

The flexibilities of the ssDNA dT-50, ssDNA dN-50, ssDNA dA-50 and the dsDNA dN-50p molecules were assessed using small-angle x-ray scattering (SAXS) at the Diamond Light Source (Harwell, UK). SAXS data were collected using a size exclusion KW 402.5 (2.4 ml) column (Shodex). 50 µl of DNA sample was injected and elution was carried out at 37°C at 75 µl/min using SPR running buffer in the absence of glycerol and BSA. The flexible cylinder model was fit to the four scattering data sets within the 0.0037 - 0.27 [1/Å] range to derive the persistence lengths. Data plotting and fitting was carried out using the SasView software for SAXS data analysis.

### Surface Plasmon Resonance

The binding kinetics of WT and F86E RAD51 on ssDNA and dsDNA were assessed *via* surface plasmon resonance (SPR) using a Biacore T200 (GE Healthcare). CM5 SPR chip flow cells were activated by injecting 100 µl of a 1:1 N-Hydroxysuccinimide (NHS), ethyl(dimethylaminopropyl) carbodiimide (EDC) mix at 10 µl/min. 100 µl of purified streptavidin (1 mg/ml) was then injected over the flow cells at 10 µl/min. Unbound amine groups were then deactivated by injecting 50 µl of 1 M ethanolamine-HCl. 50 µl of 10 mM glycine pH 2.5 was injected to remove uncoupled, sterically bound streptavidin. For all experiments, ∼ 8 - 15 RU of biotinylated DNA substrates were immobilised onto either flow cells 2, 3 or 4. After DNA immobilisation, 10 µl of purified biotin was injected over all four flow cells at 10 µl/min to block the remaining free streptavidin binding sites. Chip activation and ligand immobilisation steps were carried out at 25°C. Using high performance injections, the CM5 chip surfaces were then primed *via* 10 injections of 10 µl SPR running buffer (150 mM KCl, 20 mM Hepes pH 7.5, 2 mM DTT, 5 mg/ml BSA, 2.5 mM ATP pH 7.5, 10 mM CaCl_2_, 10 % glycerol) at 30 µl/min. WT or F86E RAD51 diluted in SPR running buffer to the specified concentration was injected at 30 µl/min (association), followed by the injection of SPR running buffer for 20 minutes at 30 µl/min (dissociation) at 37°C.

### SPR Data Processing

SPR traces were processed using the BIAEvaluation software (GE Healthcare) as following. First, the negative control flow cell trace (Fc1 or Fc3) was subtracted from the experimental flow cell trace. The trace was then vertically and horizontally aligned, such that the start of the protein injection occurs at time = 0 s, Response = 0 RU. Subsequently, the drift trace between the negative control flow cell and the experimental flow cell during the final priming injection was subtracted from the experimental flow cell trace (double-referencing) (Myszka, 1999). All experimental curves were normalised using the following equation:

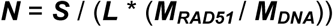

***N*** is the normalised signal (mean number of RAD51 molecules per DNA oligo), ***S*** is the signal in RU units (1 RU ∼ 50 pg/mm^2^), ***L*** is the amount of DNA ligand immobilised onto the experimental flow cell (RU), ***M_RAD51_***is the molecular weight (kDa) of RAD51 (∼ 37 kDa) and ***M_DNA_***is the molecular weight (kDa) of the immobilised DNA molecule. The normalised signal represents the mean number of RAD51 molecules bound to an immobilised DNA molecule, and has a predicted maximum at 16 (i.e. the predicted maximum number of RAD51 molecules bound to a ssDNA or dsDNA 50mer).

### Mathematical Model Development and Analysis

The ODE model describing RAD51 polymerisation on DNA consists of an equilibrium sub-model and a kinetic sub-model. The equilibrium sub-model was formulated using second order mass-action kinetics *via* the rule-based modelling language BioNetGen (Harris et al., 2016) and consists of 16 non-linear ODEs describing the formation of RAD51 polymers in solution up to a maximum length of 16 (i.e. the predicted maximum number of RAD51 molecules binding to a DNA 50-mer (Short et al., 2016)). Each ODE describes the rate of change in concentration of a RAD51 polymer in solution with respect to time. They are non-linear because monomeric RAD51 is consumed in the process of polymerisation to form RAD51 polymers. In this sub-model, we assume any RAD51 n-mer (1 ≤ n < 16) can bind to any other RAD51 m-mer (1 ≤ m < 16) to form a RAD51 (n+m)-mer (1 < n + m ≤ 16). In addition, any RAD51 k-mer can fall apart in every possible combination of m-mers and n-mers (k = n + m) (e.g. a pentamer can fall apart to form a monomer and a tetramer, or a dimer and a trimer). The single dissociation constant (K_D_) describing all of these pairwise interactions is a fit parameter derived from the SPR data. The model is evaluated at equilibrium to calculate the concentration of each RAD51 polymer in solution, prior to the SPR injection. The concentration distribution of RAD51 polymers in solution depends on the fit parameter K_D_. The predicted concentrations of RAD51 polymers are then inserted into the kinetic sub-model.

The kinetic sub-models describe the formation of RAD51 polymers on DNA. Importantly, during SPR injections the concentrations of RAD51 polymers in solution in the flow cell remains constant because RAD51 is continuously replenished by flow. For this reason, the kinetic sub-models were formulated using first order mass-action kinetics *via* the rule-based modelling language BioNetGen (Harris et al., 2016) and each consist of 17 linear ODEs describing the change in concentration of RAD51-polymer-bound DNA molecules. The extra ODE describes unbound DNA molecules. See Supplementary Information for full details of model development.

For WT RAD51, ABC-SMC was carried in MATLAB 2016b to determine a unique set of parameters for the equilibrium and kinetic sub-models that can simultaneously explain the ssDNA dN-X data (Figs. 1B, 2A, S2) and the dsDNA dN-Xp data (Figs. 1D, 2B, S2). dN-X & dN-Xp K_D_, dN-X k_p_, k_q_, k_s_, and dN-Xp k_p_ are well determined. dN-X k_u_ and dN-Xp k_u_ were undetermined, due to the fact that 1) no RAD51 binding was observed to the dN-8 and dN-5p oligos, and 2) the concentration of RAD51 monomers, dimers and trimers in solution at 3 µM [WT RAD51] is low relative to longer polymers (Fig. 2C). Finally, we could only determine a well-defined upper bound for the dsDNA stable reverse rate constant (dN-Xp k_s_ < 10^-3^). For F86E RAD51, ABC-SMC was carried out in MATLAB 2016b to determine a unique set of parameters for the equilibrium sub-model and a simplified version of the kinetic sub-model (Fig. 4D) that can simultaneously explain the ssDNA dT-50, dN-50, dA-50 and the dsDNA dN-50p data (Figs. 4C, E, H, S6). All parameters except dN-50p k_pr_ were well determined. dN-50p k_pr_ has a well defined upper bound (i.e. dN-50p k_pr_ < 10^-3^).

All data and scripts are freely available via: https://osf.io/28cqv/

### Statistical Analysis

Unpaired, Mann-Whitney-Wilcoxon tests were carried out to test for significant differences between the different fitted RAD51 polymerisation rates and lifetimes (Fig. 3 D **and** E, Fig. 4 E **and** H). All tests were carried out using GraphPad Prism 7.

## Author contributions

FE, OD and FP conceived and planned the project. FP conducted all SPR experiments, protein purification and mathematical modelling. AES and TB designed and generated the short DNA oligos for SPR experiments. JA contributed to the conceptualisation of the thermodynamic impact of polymerisation. FE, OD and FP wrote the manuscript with input from all contributing authors.

## Acknowledgements

FE and OD are supported by Wellcome Trust Senior Research Fellowships in Basic Biomedical Science (101009/Z/13/Z and 207537/Z/17/Z, respectively). JA is supported by National Science Foundation grant (DMS 1454739). TB is thankful for the support by Biotechnology and Biological Sciences Research Council (BB/J001694/2). FP is a recipient of the Systems Biology Doctoral Training Centre Scholarship, funded by the Engineering and Physical Sciences Research Council. We thank Marcus Bridge for assistance with SPR, Nicola Trendel for assistance with ABC-SMC, David Staunton for the SEC-MALS analysis of the RAD51 mutant, and Robert Rambo for the DNA oligo SAXS analysis. We also acknowledge the use of the University of Oxford Advanced Research Computing (ARC) facility in carrying out this work (http://dx.doi.org/10.5281/zenodo.22558).

## Figure & Table Legends

**Figure S1.**
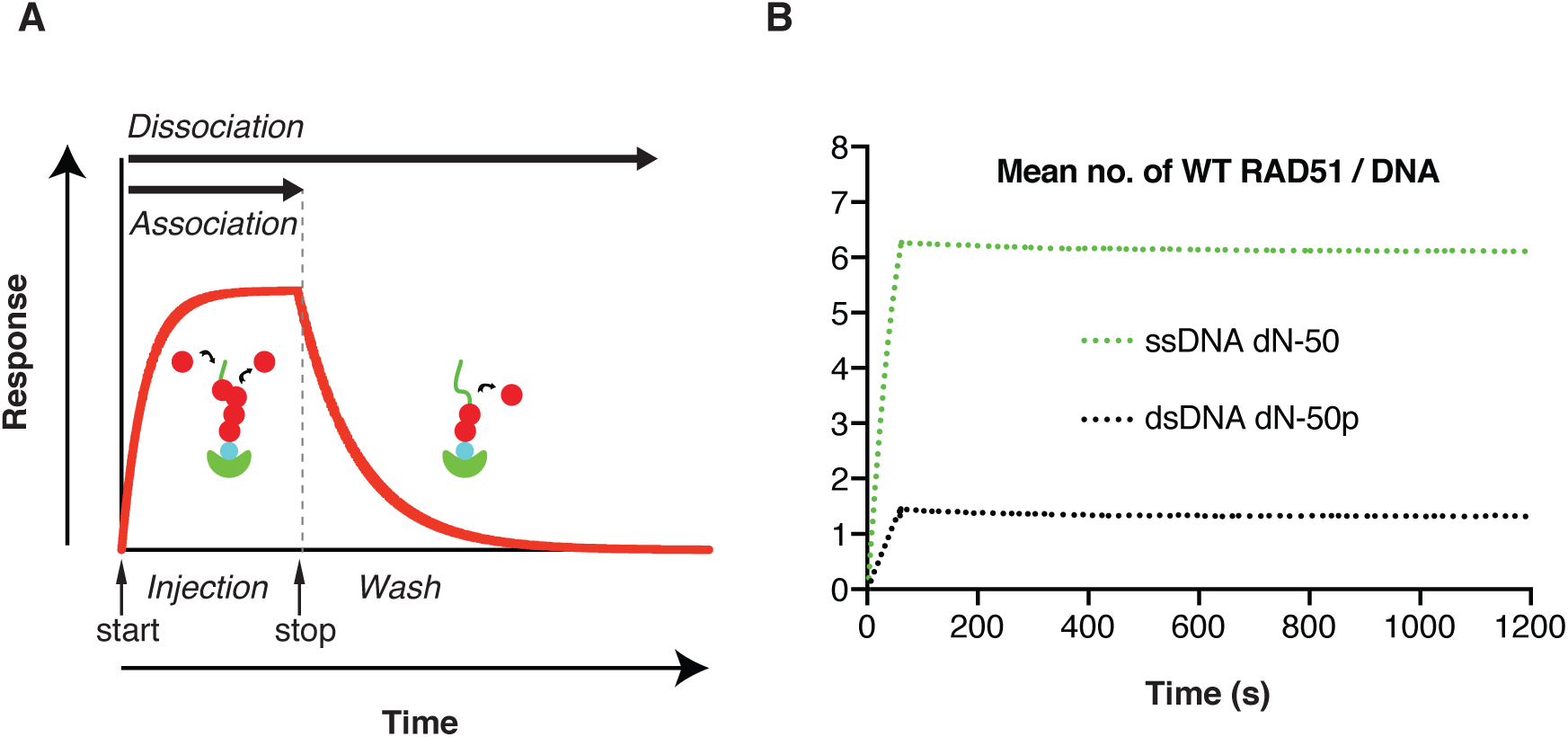
The experimental setup. ***A***. Depiction of the experimental setting. A biotinylated DNA oligo is immobilised onto a surface plasmon resonance (SPR) CM5 chip *via* biotin-streptavidin interaction. RAD51 protein is injected over the DNA-coated SPR matrix to measure polymerisation kinetics. Throughout this period, association and dissociation of RAD51 take place simultaneously. Following protein injection (stop), running buffer is injected to measure dissociation kinetics. ***B***. Wild-type (WT) RAD51 SPR curves for ssDNA dN-50 and dsDNA dN-50p. RAD51 was injected at 150 nM in the presence of 2.5 mM ATP (pH 7.5) and 10 mM CaCl_2_. Curves were normalised to the R_max_, whereby the normalised response value of X (vertical axis) indicates the average number of RAD51 molecule bound to a single DNA oligo.

**Figure S2.**
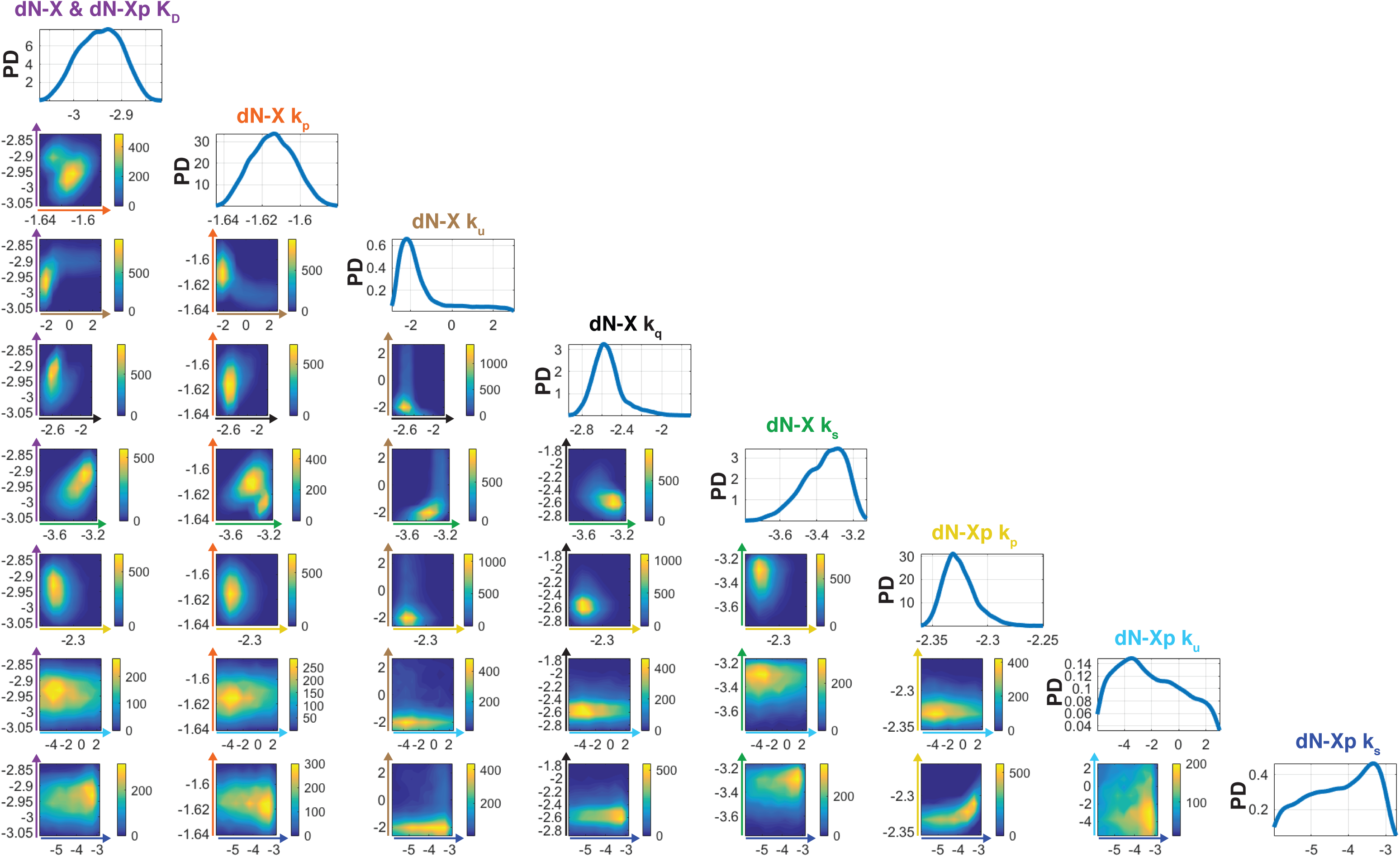
ABC-SMC for WT RAD51 SPR data fitting. ABC-SMC probability densities for the simultaneous WT RAD51 ssDNA (Fig. 2A) and dsDNA model (Fig. 2B) fits of the ssDNA dN-X (Fig. 1B) and dsDNA dN-Xp (Fig. 1D) SPR data. Heat maps indicate particle frequencies and describe any pair-wise correlations between model parameters (dN-X & dN-Xp K_D_, dN-X k_p_, k_u_, k_q_, k_s_, dN-Xp k_p_, k_u_, k_s_). All heat map axes are in log-log scale and are labelled according to the corresponding parameter colour: dN-X & dN-Xp K_D_ (purple), dN-X k_p_ (orange), dN-X k_u_ (brown), dN-X k_q_ (black), dN-X k_s_ (green), dN-Xp k_p_ (yellow), dN-Xp k_u_ (cyan), dN-Xp k_s_ (blue). A high particle frequency corresponds to a good model fit to the data for the specified parameter pair. The probability densities for individual parameters are presented along the top left diagonal and were estimated using a Gaussian kernel. Particles were initialised *via* log-uniform priors with the following lower and upper bounds: 10^-4^ < dN-X & dN-Xp K_D_ < 10^-1^, 10^-4^ < dN-X k < 10^4^, 10^-6^ < dN-X k < 10^3^, 10^-6^ < dN-X k < 10^3^, 10^-6^ < dN-X k < 10^3^, 10^-4^ < dN-Xp k < 10^4^, 10^-6^ < dN-Xp k < 10^3^, 10^-6^ < dN-Xp k < 10^3^. PD: probability density.

**Figure S3.**
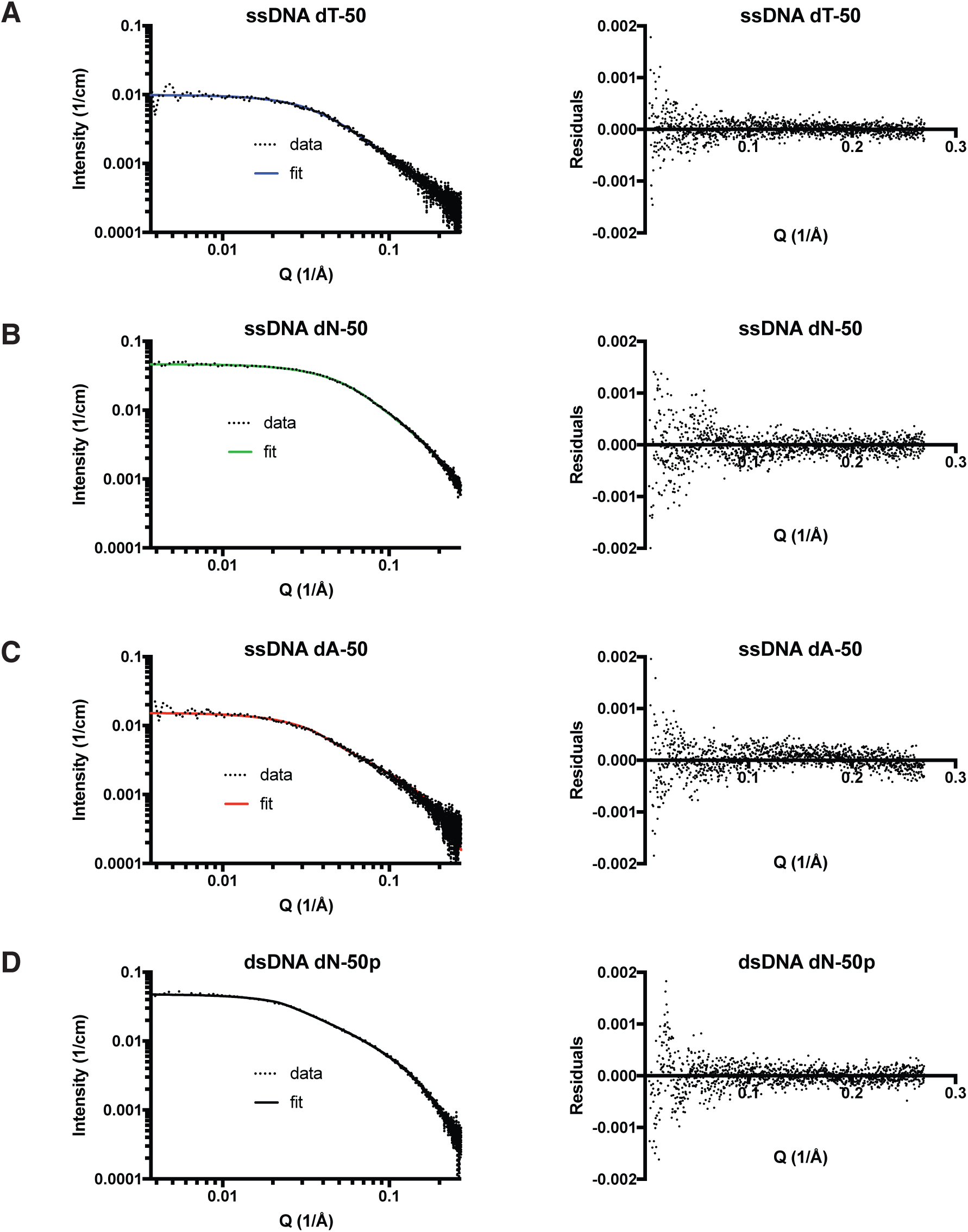
The flexibility measurements of DNA oligos used in this study. ***A-D*.** The flexibilities of ssDNA dT-50 (A), dN-50 (B), dA-50 (C) and dsDNA dN-50p (D) were assessed by small-angle x-ray scattering (SAXS). The left panels show the absolute scattering intensity (cm^-1^) measured as a function of the scattering vector (Å^-1^). The flexible cylinder model was fit to the data (fitting range: 0.0037 - 0.27 Å^-1^) to predict the kuhn lengths (persistence length = kuhn length / 2). The right panels show residuals of the flexible cylinder model fits.

**Figure S4.**
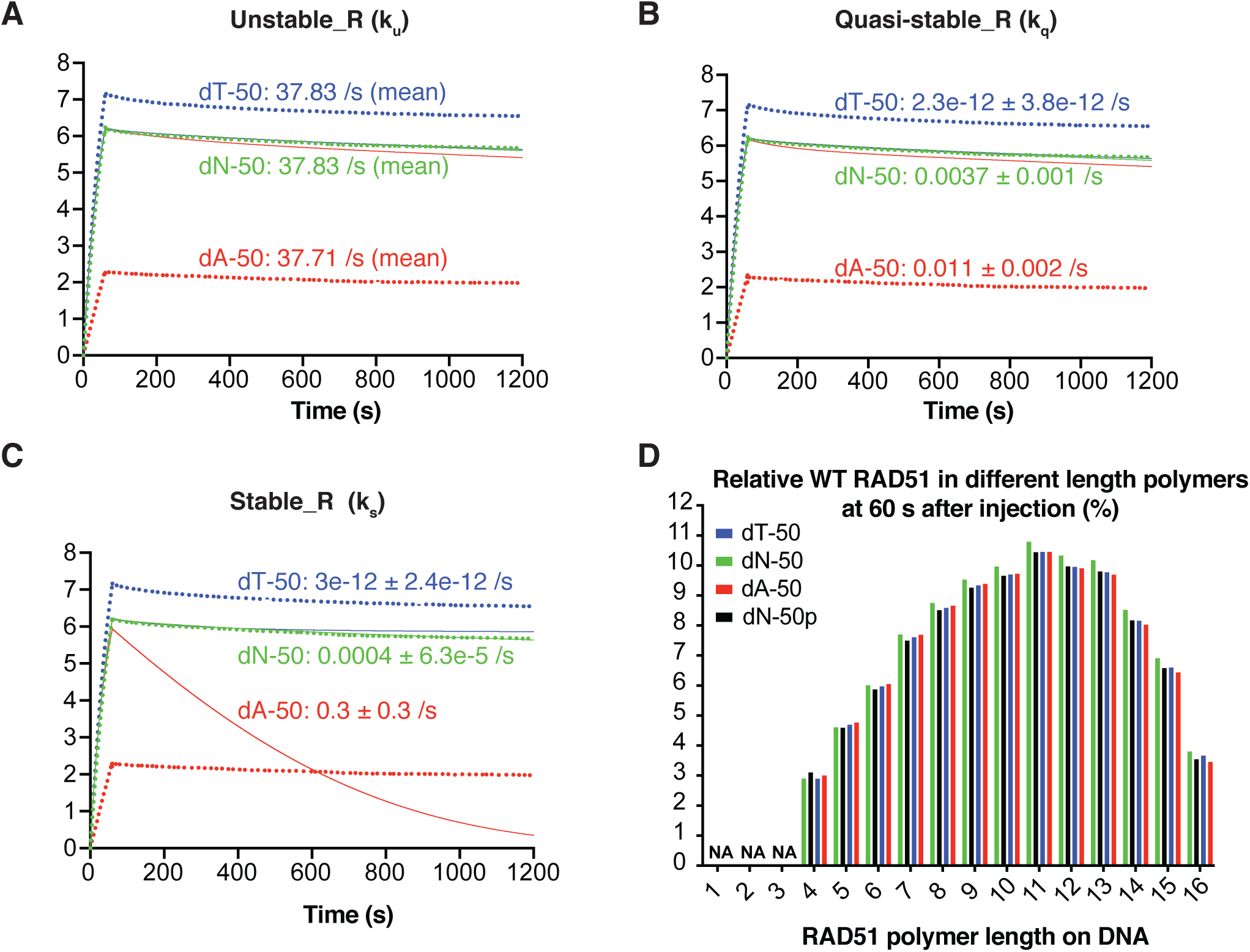
Evaluation of WT RAD51 ssDNA model for ssDNA of variable flexibility. ***A-C*.** RAD51 SPR curves (dotted lines) for the ssDNA dT-50, dN-50 and dA-50 were fitted by varying a single parameter of the ssDNA model using lsqcurvefit (MATLAB). Each panel shows the ODE model based on varying the unstable reverse rate constant (k_u_) (***A***), the quasi-stable reverse rate constant (k_q_) (***B***), and the stable reverse rate constant (k_s_) (***C***). Individually varying either k_u_, k_q_, or k_s_ does not generate accurate model fits. dN-X **k_u_** was undetermined (Fig. 2A), and for this reason fit values in A are reported only as means. ***D***. Model prediction of the % WT RAD51 in different length polymers on ssDNA dN-50, dT-50, dA-50, and dsDNA dN-50p after a 60 second injection of WT RAD51 using models from Fig. 2 A **and** B. Parameter values in the ssDNA model were set to the mean values as identified in Fig. 2A, and % WT RAD51 values were calculated by multiplying the polymer concentrations by their respective polymer length and dividing each value by the total amount of RAD51 bound to DNA after a 60 second injection (i.e. % WT RAD51 = [*n*-mer] * *n* / [WT RAD51_total DNA-bound_]). dN-X and dN-Xp **k_u_** were undetermined (Fig. 2 A **and** B), and for this reason the values for % WT RAD51 in monomers, dimers, and trimers were not reported and labelled as NA (not-applicable).

**Figure S5.**
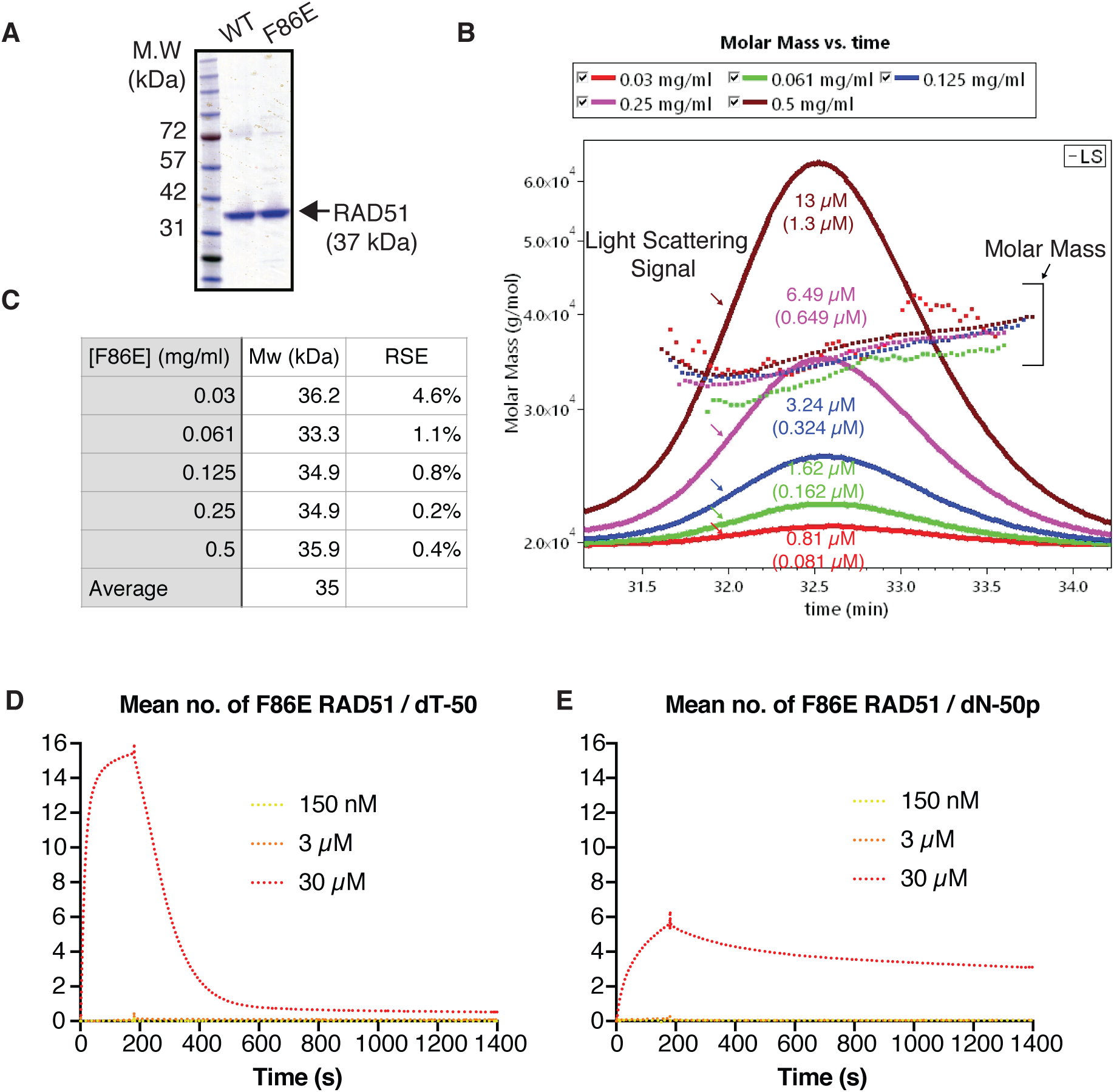
Biochemical characterisation of F86E RAD51 used in this study. ***A.*** Purified WT and F86E RAD51 on an SDS-PAGE gel are visualized by coomassie blue staining. ***B.*** Size-exclusion multi-angle light scattering (SEC-MALS) curves for F86E RAD51 at five injection concentrations as indicated. The Superdex200 10/300 GL size exclusion column leads to a ∼ 10-fold dilution of the injected sample. Hence, approximate elution concentrations are given in brackets below the injection concentrations. Dotted lines represent the molar mass derived values for each elution peak, solid lines represent light scattering. F86E RAD51 concentrations are shown in both mg/ml and µM. ***C*.** Calculated molecular weights of the five F86E RAD51 elution peaks. Data were analysed using the ASTRA 6.1 chromatography software for SEC-MALS. RSE: relative standard error. ***D* and *E*.** F86E RAD51 displays reduced affinity for ssDNA and dsDNA. F86E RAD51 SPR binding curves for the ssDNA dT-50 (D) and the dsDNA dN-50p (E) at the colour-coded concentrations are shown.

**Figure S6.**
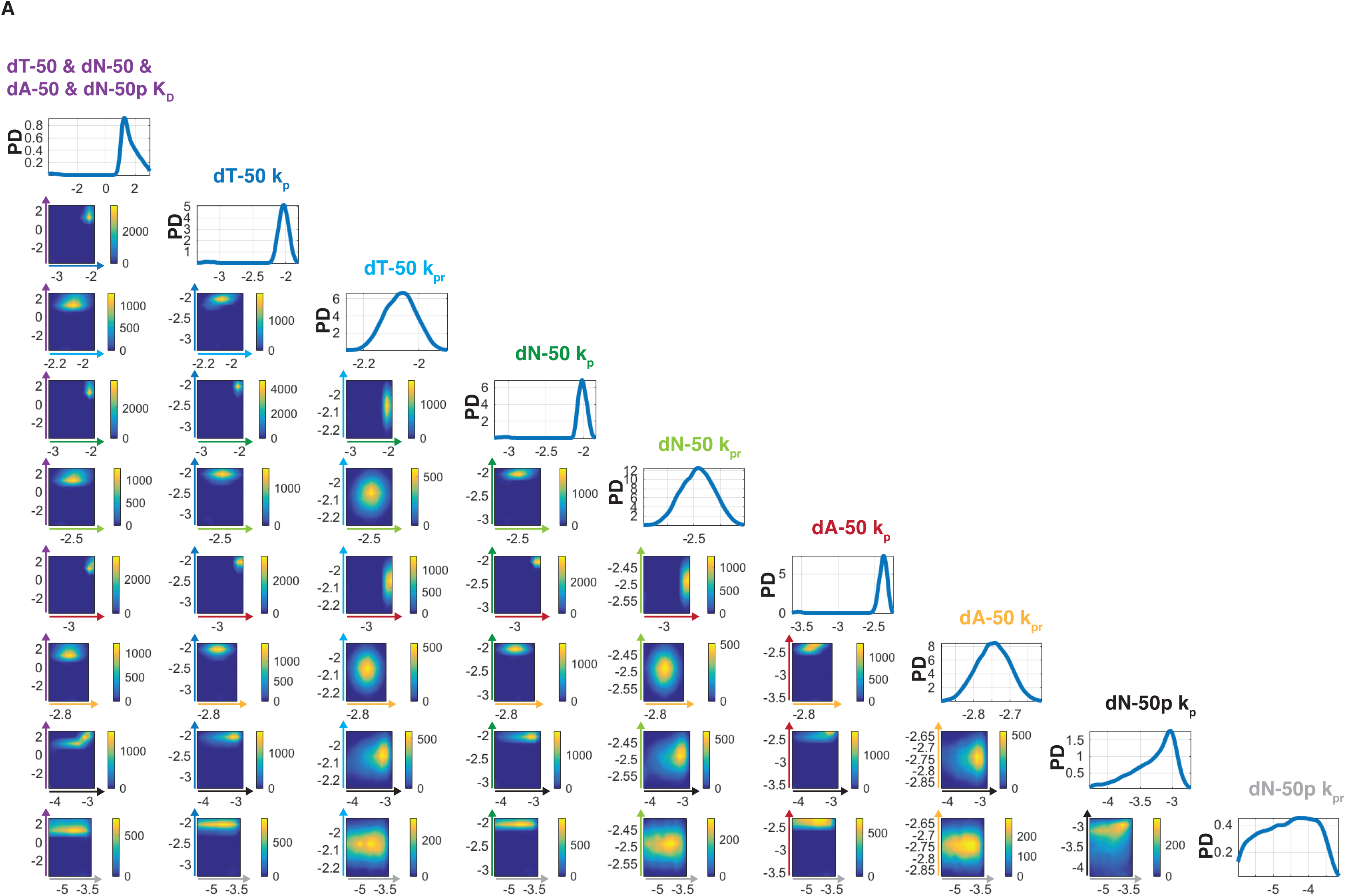
ABC-SMC for F86E RAD51 kinetic data fitting. ABC-SMC probability densities for the F86E RAD51 ssDNA/dsDNA model (Fig. 4D) fit of the ssDNA dN-50, dA-50, dT-50 and dsDNA dN-50p SPR data. All four SPR curves were fit simultaneously using the same model. k_p_ and k_pr_ values were individually fit to each curve. A single K_D_ was fit to all four curves. Heat maps indicate particle frequencies and describe any pair-wise correlations between model parameters (dT-50 & dN-50 & dA-50 & dN-50p K_D_, dT-50 k_p_, k_pr_, dN-50 k_p_, k_pr_, dA-50 k_p_, k_pr_, dN-50p k_p_, k_pr_). All heat map axes are in log-log scale and are labelled according to the corresponding parameter colour: dN-X & dN-Xp K_D_ (purple), dT-50 k_p_ (blue), dT-50 k_pr_ (cyan), dN-50 k_p_ (green), dN-50 k_pr_ (light green), dA-50 k_p_ (red), dA-50 k_pr_ (salmon), dN-50p k_p_ (black), dN-50p k_pr_ (grey). A high particle frequency corresponds to a good model fit to the data for the specified parameter pair. The probability densities for individual parameters are presented along the top left diagonal and were estimated using a Gaussian kernel. Particles were initialised *via* log-uniform priors with the following lower and upper bounds: 10^-4^ < K_D_ < 10^3^, 10^-6^ < dT-50 k_p_ < 10^4^, 10^-6^ < dT-50 k_pr_ < 10^3^, 10^-6^ < dN-50 k_p_ < 10^4^, 10^-6^ < dN-50 k_pr_ < 10^3^, 10^-6^ < dA-50 k_p_ < 10^4^, 10^-6^ < dA-50 k_pr_ < 10^3^, 10^-6^ < dN-50p k_p_ < 10^4^, 10^-6^ < dN-50p k_pr_ < 10^3^. PD: probability density.

