## Supporting information for "Molecular flexibility of DNA as a key determinant of RAD51 recruitment"

Federico Paoletti[1], Afaf El-Sagheer[2,3], Jun Allard[4],

Tom Brown[2], Omer Dushek[1,†], Fumiko Esashi[1,†]

1. Sir William Dunn School of Pathology, University of Oxford, OX1 3RE, UK.

2. Department of Chemistry, University of Oxford, OX1 3TA, UK.

3. Department of Science and Mathematics, Suez University, Suez 43721, Egypt.

4. Department of Mathematics, University of California, Irvine, CA 92697-4575, US.

[†]Equal contribution and corresponding authors:

Omer Dushek:

Fumiko Esashi:

### 1 Mathematical Model Development and Analysis

The ssDNA and dsDNA models describing RAD51 polymerisation on ssDNA and dsDNA, referred to as the $n$-mer models, each consist of an equilibrium sub-model and a kinetic sub-model. The equilibrium sub-model was formulated using second order mass-action kinetics via the rule-based modelling language BioNetGen [4], and consists of 16 non-linear ODEs describing the formation of RAD51 polymers in solution up to a maximum length of 16 (i.e. the predicted maximum number of RAD51 molecules binding to a DNA 50-mer [6]). Each ODE describes the rate of change in concentration of a RAD51 polymer in solution with respect to time. They are non-linear because monomeric RAD51 is consumed in the process of polymerisation to form RAD51 polymers. In this sub-model, any RAD51 $n$-mer ($1 \leq n < 16$) can bind to any other RAD51 $m$-mer ($1 \leq m < 16$) to form a RAD51 $(n+m)$-mer ($1 < n+m \leq 16$). In addition, any RAD51 $k$-mer can fall apart in every possible combination of $m$-mers and $n$-mers ($k = n + m$) (e.g. a pentamer can fall apart to form a monomer and a tetramer, or a dimer and a trimer). The single dissociation constant ($\mathbf{K_D}$) describing all of these pairwise interactions is a fit parameter derived from the SPR data. The model is evaluated at equilibrium to calculate the concentration of each RAD51 polymer in solution, prior to the SPR injection. The concentration distribution of RAD51 polymers in solution depends on the fit parameter $\mathbf{K_D}$ and on the RAD51 monomer concentration (i.e. the RAD51 SPR injection concentration). The predicted concentrations of RAD51 polymers are then inserted into the kinetic sub-models.

The ssDNA and dsDNA kinetic sub-models describe the formation of RAD51 polymers on ssDNA and dsDNA, respectively. Importantly, during SPR injections the concentrations of RAD51 polymers in solution in the flow cell remains constant because

RAD51 is continuously replenished by flow. For this reason, the kinetic sub-models were formulated using first order mass-action kinetics via the rule-based modelling language BioNetGen [4] and each consist of 17 linear ODEs. Each ODE describes the rate of change in mean concentration of a DNA-bound RAD51 $n$-mer on the SPR chip with respect to time, where $1 \leq n \leq b$ and $b$ is the maximum number of RAD51 binding sites on the DNA molecule (Table 1).

All data and scripts are freely available here.

Table 1: DNA molecule names and lengths, with corresponding maximum RAD51 polymer lengths ($b$) and n-mer model (NM) ODEs used for SPR data fitting. SPR data obtained using the ssDNA dN-5 and dN-11 oligos was not included in the data fitting because they were identical to the data obtained using the ssDNA dN-8 oligo and described no RAD51 binding to these oligos. NA: not applicable. ESM: equilibrium sub-model.

| Molecule Name | Length (nt or bp) | $b$ | NM ODEs |
|---|---|---|---|
| ssDNA dN-5 | 5 | 1 | NA |
| dsDNA dN-5p | 5 | 1 | ESM, 10 - 11 |
| ssDNA dN-8 | 8 | 2 | ESM, 5 - 7 |
| dsDNA dN-8p | 8 | 2 | ESM, 10 - 12 |
| ssDNA dN-11 | 11 | 3 | NA |
| dsDNA dN-11p | 11 | 3 | ESM, 10 - 13 |
| ssDNA dN-14 | 14 | 4 | ESM, 5 - 8 |
| ssDNA dN-17 | 17 | 5 | ESM, 5 - 8 |
| ssDNA dN-50 | 50 | 16 | ESM, 5 - 9 |
| ssDNA dT-50 | 50 | 16 | ESM, 5 - 9 |
| ssDNA dA-50 | 50 | 16 | ESM, 5 - 9 |
| dsDNA dN-50p | 50 | 16 | ESM, 10 - 14 |

#### 1.1 Assumptions

For the equilibrium sub-model, we assume

- any RAD51 $n$-mer ($1 \leq n < 16$) can bind to any other RAD51 $m$-mer ($1 \leq m < 16$) to form a RAD51 $(n+m)$-mer ($1 < n+m \leq 16$)
- any RAD51 $k$-mer can fall apart in every possible combination of $m$-mers and $n$-mers ($k = n + m$)
- the dissociation constant between RAD51 polymers $\mathbf{K_D}$ is independent of RAD51 polymer length: $\mathbf{K_D}(n, m) = \mathbf{K_D}(1, 1) = \mathbf{K_D}$. This assumption was previously applied in [2]

For the ssDNA and dsDNA kinetic sub-models, we assume

- Each ssDNA or dsDNA molecule can accommodate a RAD51 polymer of maximum length $b$, where $b$ is the maximum number of RAD51 binding sites on the DNA molecule (Table 1)
- RAD51 only nucleates once per DNA molecule (i.e. each DNA molecule only accommodates a single polymer of length $n$, where $1 \leq n \leq b$). In other words, polymer elongation is faster than nucleation. This is a valid assumption given that the longest ssDNA and dsDNA molecules used in this study are 50 nt and 50 bp long respectively, and RAD51 nucleation was shown to occur on average once every 500 bp in reactions with a 1 RAD51 : 3 bp molar ratio [5]. In the case of the experiments using WT RAD51 and the ssDNA dN-50 or dsDNA dN-50p, we used a 150 nM RAD51 concentration. Considering an average 10 RU of immobilised dsDNA and a dN-50p molecular weight of 31.157 kDa, the molar ratio in my experimental system is far lower compared to that in [5],

and hence nucleation is expected on average to occur only once on a DNA molecule: [bps] $= 50(\frac{\text{R}_{\text{lig}}}{100\text{M}_{\text{lig}}})[3] = 50(\frac{10\text{RU}}{(100)31157\text{Da}}) \approx 160\,\mu$M. Hence $\frac{150\text{nM RAD51}}{160\mu\text{M bps}} \Rightarrow$ 1 RAD51 molecule per $\approx$ 1100 bps on the SPR chip.

- Any RAD51 $n$-mer ($1 \leq n \leq b$) can adsorb onto DNA from solution. If $n < b$, further RAD51 polymers in solution can elongate the DNA-bound $n$-mer to form a DNA-bound ($n^*$)-mer ($1 < n^* \leq b$)

- The rate of adsorption of RAD51 $n$-mers on DNA is the same as the rate of elongation of RAD51 $n$-mers on DNA. This rate is described as the polymerisation rate constant $\mathbf{k_p}$

- RAD51 dissociation occurs via the removal of single RAD51 protomers and of short, unstable RAD51 nuclei. This is a valid assumption when considering that in all the SPR experiments in this study ATP hydrolysis is inhibited via the addition of $Ca^{2+}$, leading to very slow RAD51 filament disassembly which can be approximated as disassembly via monomer removal or via the removal of short, unstable RAD51 nuclei

#### 1.2 Description of ODEs - Equilibrium Sub-Model

A previous study applied a closed form expression to evaluate the concentration of RAD51 $n$-mers as a function of $\mathbf{K_D}$ and the RAD51 monomer concentration [2], but did not provide a derivation of the expression. For this reason, we preferred to develop a comprehensive ODE model using BioNetGen [4] that takes account of all $n$-mer + $m$-mer interactions. The BioNetGen code for the equilibrium sub-model is as follows:

```
begin parameters

ka
kd
Rad51m

end parameters

begin molecule types

R_1()
R_2()
R_3()
R_4()
R_5()
R_6()
R_7()
R_8()
R_9()
R_10()
R_11()
R_12()
R_13()
R_14()
R_15()
R_16()

end molecule types

begin seed species

R_1()        Rad51m

end seed species

begin reaction rules

R_1() + R_1() <-> R_2()                ka, kd
R_1() + R_2() <-> R_3()                ka, kd
R_1() + R_3() <-> R_4()                ka, kd
R_1() + R_4() <-> R_5()                ka, kd
R_1() + R_5() <-> R_6()                ka, kd
R_1() + R_6() <-> R_7()                ka, kd
```

```
R_1() + R_7() <-> R_8()                ka, kd
R_1() + R_8() <-> R_9()                ka, kd
R_1() + R_9() <-> R_10()                ka, kd
R_1() + R_10() <-> R_11()                ka, kd
R_1() + R_11() <-> R_12()                ka, kd
R_1() + R_12() <-> R_13()                ka, kd
R_1() + R_13() <-> R_14()                ka, kd
R_1() + R_14() <-> R_15()                ka, kd
R_1() + R_15() <-> R_16()                ka, kd

R_2() + R_2() <-> R_4()                ka, kd
R_2() + R_3() <-> R_5()                ka, kd
R_2() + R_4() <-> R_6()                ka, kd
R_2() + R_5() <-> R_7()                ka, kd
R_2() + R_6() <-> R_8()                ka, kd
R_2() + R_7() <-> R_9()                ka, kd
R_2() + R_8() <-> R_10()                ka, kd
R_2() + R_9() <-> R_11()                ka, kd
R_2() + R_10() <-> R_12()                ka, kd
R_2() + R_11() <-> R_13()                ka, kd
R_2() + R_12() <-> R_14()                ka, kd
R_2() + R_13() <-> R_15()                ka, kd
R_2() + R_14() <-> R_16()                ka, kd

R_3() + R_3() <-> R_6()                ka, kd
R_3() + R_4() <-> R_7()                ka, kd
R_3() + R_5() <-> R_8()                ka, kd
R_3() + R_6() <-> R_9()                ka, kd
R_3() + R_7() <-> R_10()                ka, kd
R_3() + R_8() <-> R_11()                ka, kd
R_3() + R_9() <-> R_12()                ka, kd
R_3() + R_10() <-> R_13()                ka, kd
R_3() + R_11() <-> R_14()                ka, kd
R_3() + R_12() <-> R_15()                ka, kd
R_3() + R_13() <-> R_16()                ka, kd

R_4() + R_4() <-> R_8()                ka, kd
R_4() + R_5() <-> R_9()                ka, kd
R_4() + R_6() <-> R_10()                ka, kd
R_4() + R_7() <-> R_11()                ka, kd
R_4() + R_8() <-> R_12()                ka, kd
R_4() + R_9() <-> R_13()                ka, kd
R_4() + R_10() <-> R_14()                ka, kd
```

```
R_4() + R_11() <-> R_15()              ka, kd
R_4() + R_12() <-> R_16()              ka, kd

R_5() + R_5() <-> R_10()              ka, kd
R_5() + R_6() <-> R_11()              ka, kd
R_5() + R_7() <-> R_12()              ka, kd
R_5() + R_8() <-> R_13()              ka, kd
R_5() + R_9() <-> R_14()              ka, kd
R_5() + R_10() <-> R_15()              ka, kd
R_5() + R_11() <-> R_16()              ka, kd

R_6() + R_6() <-> R_12()              ka, kd
R_6() + R_7() <-> R_13()              ka, kd
R_6() + R_8() <-> R_14()              ka, kd
R_6() + R_9() <-> R_15()              ka, kd
R_6() + R_10() <-> R_16()              ka, kd

R_7() + R_7() <-> R_14()              ka, kd
R_7() + R_8() <-> R_15()              ka, kd
R_7() + R_9() <-> R_16()              ka, kd

R_8() + R_8() <-> R_16()              ka, kd

end reaction rules

begin observables

Molecules Rad51_16mers R_16()
Molecules Rad51_15mers R_15()
Molecules Rad51_14mers R_14()
Molecules Rad51_13mers R_13()
Molecules Rad51_12mers R_12()
Molecules Rad51_11mers R_11()
Molecules Rad51_10mers R_10()
Molecules Rad51_9mers R_9()
Molecules Rad51_Octamers R_8()
Molecules Rad51_Heptamers R_7()
Molecules Rad51_Hexamers R_6()
Molecules Rad51_Pentamers R_5()
Molecules Rad51_Tetramers R_4()
Molecules Rad51_Trimers R_3()
Molecules Rad51_Dimers R_2()
Molecules Rad51_Monomers R_1()
```

```
end observables

generate_network({overwrite=>1})
writeMfile({})
```

In the first section (`begin parameters`) all model parameters, including kinetic rate constants, are defined. In the second section (`begin molecule types`), all possible RAD51 polymer lengths in solution are defined. In the third section (`begin seed species`), the initial condition for $R_1$ is defined ($R_1(0) =$`Rad51m`$=$ RAD51 injection concentration). In the fourth section (`begin reaction rules`), all allowed RAD51 polymerisation reactions in solution are defined with their associated kinetic rate constants. The final section (`begin observables`) defines the desired model outputs (in this case $nR_n$ for $n = 1, ..., 16$).

The resulting ODEs for the equilibrium sub-model in compact form are as follows:

for $n = 1$:

$$\frac{dR_1}{dt} = k_\mathsf{d}(\frac{1}{2}R_2 + \sum_{3}^{16} R_i) - k_\mathsf{a}R_1(2R_1 + \sum_{2}^{15} R_i) \tag{1}$$

for $2 \leq n \leq 8$:

$$\frac{dR_n}{dt} = k_\mathsf{d}(\sum_{n+1}^{2n-1} R_i + \frac{1}{2}R_{2n} + \sum_{2n+1}^{16} R_i - \left\lfloor\frac{n}{2}\right\rfloor R_n) - k_\mathsf{a}[R_n(\sum_{1}^{n-1} R_i + 2R_n + \sum_{n+1}^{16-n} R_i) - \sum_{1}^{\lfloor\frac{n}{2}\rfloor} R_i R_{n-i}] \tag{2}$$

for $9 \leq n \leq 15$:

$$\frac{dR_n}{dt} = k_\mathsf{d}(\sum_{n+1}^{16} R_i - \left\lfloor\frac{n}{2}\right\rfloor R_n) - k_\mathsf{a}(R_n \sum_{1}^{16-n} R_i - \sum_{1}^{\lfloor\frac{n}{2}\rfloor} R_i R_{n-i}) \tag{3}$$

for $n = 16$:

$$\frac{dR_{16}}{dt} = k_\mathsf{a} \sum_{1}^{8} R_i R_{16-i} - 8k_\mathsf{d}R_{16} \tag{4}$$

with initial conditions $R_1(0) =$ RAD51 SPR injection concentration, $R_{2,\ldots,16}(0) = 0$.

$R_n$ is the time-dependent concentration of RAD51 $n$-mers in solution, $k_\mathsf{a}$ and $k_\mathsf{d}$ are the association and dissociation rate constants of the RAD51 protomer-protomer interaction, and $\lfloor . \rfloor$ is the floor function (e.g. $\lfloor 2.7 \rfloor = 2$). To fit the $\mathbf{K_D}$ of the RAD51 protomer-protomer interaction, $k_\mathsf{a}$ was set to 1 and the $k_\mathsf{d}$ was fit by simulating the model at equilibrium. Hence, $\mathbf{K_D} = \frac{k_d}{k_a} = k_d$.

#### 1.3 Description of ODEs - Kinetic Sub-Models

The ODEs for the general ssDNA and dsDNA kinetic sub-models were both generated via the same BioNetGen code:

```
begin parameters

###CONCENTRATIONS FROM RAD51 EQUILIBRIUM SUB-MODEL
Rad51_monomer
Rad51_dimer
Rad51_trimer
Rad51_tetramer
Rad51_pentamer
Rad51_hexamer
Rad51_heptamer
Rad51_8mer
Rad51_9mer
Rad51_10mer
Rad51_11mer
Rad51_12mer
Rad51_13mer
Rad51_14mer
Rad51_15mer
Rad51_16mer
##################################################

DNA_start

kp

kd_1
kd_2
kd_3
kd_4
kd_5
```

```
kd_6
kd_7
kd_8
kd_9
kd_10
kd_11
kd_12
kd_13
kd_14
kd_15
kd_16

ks

kadf_obs_1        kp * Rad51_monomer
kadf_obs_2        kp * Rad51_dimer
kadf_obs_3        kp * Rad51_trimer
kadf_obs_4        kp * Rad51_tetramer
kadf_obs_5        kp * Rad51_pentamer
kadf_obs_6        kp * Rad51_hexamer
kadf_obs_7        kp * Rad51_heptamer
kadf_obs_8        kp * Rad51_8mer
kadf_obs_9        kp * Rad51_9mer
kadf_obs_10        kp * Rad51_10mer
kadf_obs_11        kp * Rad51_11mer
kadf_obs_12        kp * Rad51_12mer
kadf_obs_13        kp * Rad51_13mer
kadf_obs_14        kp * Rad51_14mer
kadf_obs_15        kp * Rad51_15mer
kadf_obs_16        kp * Rad51_16mer

kelf_obs_1        kp * Rad51_monomer
kelf_obs_2        kp * Rad51_dimer
kelf_obs_3        kp * Rad51_trimer
kelf_obs_4        kp * Rad51_tetramer
kelf_obs_5        kp * Rad51_pentamer
kelf_obs_6        kp * Rad51_hexamer
kelf_obs_7        kp * Rad51_heptamer
kelf_obs_8        kp * Rad51_8mer
kelf_obs_9        kp * Rad51_9mer
kelf_obs_10        kp * Rad51_10mer
kelf_obs_11        kp * Rad51_11mer
kelf_obs_12        kp * Rad51_12mer
```

```
kelf_obs_13        kp * Rad51_13mer
kelf_obs_14        kp * Rad51_14mer
kelf_obs_15        kp * Rad51_15mer


end parameters

begin molecule types

DNA_0()
DNA_1()
DNA_2()
DNA_3()
DNA_4()
DNA_5()
DNA_6()
DNA_7()
DNA_8()
DNA_9()
DNA_10()
DNA_11()
DNA_12()
DNA_13()
DNA_14()
DNA_15()
DNA_16()

end molecule types

begin seed species

DNA_0()        DNA_start

end seed species

begin reaction rules

#ADSORPTION REACTIONS

DNA_0()        <-> DNA_1()                kadf_obs_1, kd_1
DNA_0() <-> DNA_2()                kadf_obs_2, kd_2
DNA_0() <-> DNA_3()                kadf_obs_3, kd_3
DNA_0() <-> DNA_4()                kadf_obs_4, kd_4
```

```
DNA_0() <-> DNA_5()                kadf_obs_5, kd_5
DNA_0() <-> DNA_6()                kadf_obs_6, kd_6
DNA_0() <-> DNA_7()                kadf_obs_7, kd_7
DNA_0() <-> DNA_8()                kadf_obs_8, kd_8
DNA_0() <-> DNA_9()                kadf_obs_9, kd_9
DNA_0() <-> DNA_10()                kadf_obs_10, kd_10
DNA_0() <-> DNA_11()                kadf_obs_11, kd_11
DNA_0() <-> DNA_12()                kadf_obs_12, kd_12
DNA_0() <-> DNA_13()                kadf_obs_13, kd_13
DNA_0() <-> DNA_14()                kadf_obs_14, kd_14
DNA_0() <-> DNA_15()                kadf_obs_15, kd_15
DNA_0() <-> DNA_16()                kadf_obs_16, kd_16

#MONOMER ELONGATION REACTIONS

DNA_1()        <-> DNA_2()                kelf_obs_1, ks
DNA_2()        <-> DNA_3()                kelf_obs_1, ks
DNA_3()        <-> DNA_4()                kelf_obs_1, ks
DNA_4() <-> DNA_5()                kelf_obs_1, ks
DNA_5() <-> DNA_6()                kelf_obs_1, ks
DNA_6() <-> DNA_7()                kelf_obs_1, ks
DNA_7() <-> DNA_8()                kelf_obs_1, ks
DNA_8() <-> DNA_9()                kelf_obs_1, ks
DNA_9() <-> DNA_10()                kelf_obs_1, ks
DNA_10() <-> DNA_11()                kelf_obs_1, ks
DNA_11() <-> DNA_12()                kelf_obs_1, ks
DNA_12() <-> DNA_13()                kelf_obs_1, ks
DNA_13() <-> DNA_14()                kelf_obs_1, ks
DNA_14() <-> DNA_15()                kelf_obs_1, ks
DNA_15() <-> DNA_16()                kelf_obs_1, ks

#DIMER ELONGATION REACTIONS

DNA_1() -> DNA_3()                kelf_obs_2
DNA_2() -> DNA_4()                kelf_obs_2
DNA_3() -> DNA_5()                kelf_obs_2
DNA_4() -> DNA_6()                kelf_obs_2
DNA_5() -> DNA_7()                kelf_obs_2
DNA_6() -> DNA_8()                kelf_obs_2
DNA_7() -> DNA_9()                kelf_obs_2
DNA_8() -> DNA_10()                kelf_obs_2
DNA_9() -> DNA_11()                kelf_obs_2
DNA_10() -> DNA_12()                kelf_obs_2
```

```
DNA_11() -> DNA_13()                kelf_obs_2
DNA_12() -> DNA_14()                kelf_obs_2
DNA_13() -> DNA_15()                kelf_obs_2
DNA_14() -> DNA_16()                kelf_obs_2

#TRIMER ELONGATION REACTIONS

DNA_1() -> DNA_4()                kelf_obs_3
DNA_2() -> DNA_5()                kelf_obs_3
DNA_3() -> DNA_6()                kelf_obs_3
DNA_4() -> DNA_7()                kelf_obs_3
DNA_5() -> DNA_8()                kelf_obs_3
DNA_6() -> DNA_9()                kelf_obs_3
DNA_7() -> DNA_10()                kelf_obs_3
DNA_8() -> DNA_11()                kelf_obs_3
DNA_9() -> DNA_12()                kelf_obs_3
DNA_10() -> DNA_13()                kelf_obs_3
DNA_11() -> DNA_14()                kelf_obs_3
DNA_12() -> DNA_15()                kelf_obs_3
DNA_13() -> DNA_16()                kelf_obs_3

#TETRAMER ELONGATION REACTIONS

DNA_1() -> DNA_5()                kelf_obs_4
DNA_2() -> DNA_6()                kelf_obs_4
DNA_3() -> DNA_7()                kelf_obs_4
DNA_4() -> DNA_8()                kelf_obs_4
DNA_5() -> DNA_9()                kelf_obs_4
DNA_6() -> DNA_10()                kelf_obs_4
DNA_7() -> DNA_11()                kelf_obs_4
DNA_8() -> DNA_12()                kelf_obs_4
DNA_9() -> DNA_13()                kelf_obs_4
DNA_10() -> DNA_14()                kelf_obs_4
DNA_11() -> DNA_15()                kelf_obs_4
DNA_12() -> DNA_16()                kelf_obs_4

#PENTAMER ELONGATION REACTIONS

DNA_1() -> DNA_6()                kelf_obs_5
DNA_2() -> DNA_7()                kelf_obs_5
DNA_3() -> DNA_8()                kelf_obs_5
DNA_4() -> DNA_9()                kelf_obs_5
DNA_5() -> DNA_10()                kelf_obs_5
```

```
DNA_6() -> DNA_11()                kelf_obs_5
DNA_7() -> DNA_12()                kelf_obs_5
DNA_8() -> DNA_13()                kelf_obs_5
DNA_9() -> DNA_14()                kelf_obs_5
DNA_10() -> DNA_15()                kelf_obs_5
DNA_11() -> DNA_16()                kelf_obs_5

#HEXAMER ELONGATION REACTIONS

DNA_1() -> DNA_7()                kelf_obs_6
DNA_2() -> DNA_8()                kelf_obs_6
DNA_3() -> DNA_9()                kelf_obs_6
DNA_4() -> DNA_10()                kelf_obs_6
DNA_5() -> DNA_11()                kelf_obs_6
DNA_6() -> DNA_12()                kelf_obs_6
DNA_7() -> DNA_13()                kelf_obs_6
DNA_8() -> DNA_14()                kelf_obs_6
DNA_9() -> DNA_15()                kelf_obs_6
DNA_10() -> DNA_16()                kelf_obs_6

#HEPTAMER ELONGATION REACTIONS

DNA_1() -> DNA_8()                kelf_obs_7
DNA_2() -> DNA_9()                kelf_obs_7
DNA_3() -> DNA_10()                kelf_obs_7
DNA_4() -> DNA_11()                kelf_obs_7
DNA_5() -> DNA_12()                kelf_obs_7
DNA_6() -> DNA_13()                kelf_obs_7
DNA_7() -> DNA_14()                kelf_obs_7
DNA_8() -> DNA_15()                kelf_obs_7
DNA_9() -> DNA_16()                kelf_obs_7

#8-MER ELONGATION REACTIONS

DNA_1() -> DNA_9()                kelf_obs_8
DNA_2() -> DNA_10()                kelf_obs_8
DNA_3() -> DNA_11()                kelf_obs_8
DNA_4() -> DNA_12()                kelf_obs_8
DNA_5() -> DNA_13()                kelf_obs_8
DNA_6() -> DNA_14()                kelf_obs_8
DNA_7() -> DNA_15()                kelf_obs_8
DNA_8() -> DNA_16()                kelf_obs_8
```

```
#9-MER ELONGATION REACTIONS

DNA_1() -> DNA_10()                kelf_obs_9
DNA_2() -> DNA_11()                kelf_obs_9
DNA_3() -> DNA_12()                kelf_obs_9
DNA_4() -> DNA_13()                kelf_obs_9
DNA_5() -> DNA_14()                kelf_obs_9
DNA_6() -> DNA_15()                kelf_obs_9
DNA_7() -> DNA_16()                kelf_obs_9

#10-MER ELONGATION REACTIONS

DNA_1() -> DNA_11()                kelf_obs_10
DNA_2() -> DNA_12()                kelf_obs_10
DNA_3() -> DNA_13()                kelf_obs_10
DNA_4() -> DNA_14()                kelf_obs_10
DNA_5() -> DNA_15()                kelf_obs_10
DNA_6() -> DNA_16()                kelf_obs_10

#11-MER ELONGATION REACTIONS

DNA_1() -> DNA_12()                kelf_obs_11
DNA_2() -> DNA_13()                kelf_obs_11
DNA_3() -> DNA_14()                kelf_obs_11
DNA_4() -> DNA_15()                kelf_obs_11
DNA_5() -> DNA_16()                kelf_obs_11

#12-MER ELONGATION REACTIONS

DNA_1() -> DNA_13()                kelf_obs_12
DNA_2() -> DNA_14()                kelf_obs_12
DNA_3() -> DNA_15()                kelf_obs_12
DNA_4() -> DNA_16()                kelf_obs_12

#13-MER ELONGATION REACTIONS

DNA_1() -> DNA_14()                kelf_obs_13
DNA_2() -> DNA_15()                kelf_obs_13
DNA_3() -> DNA_16()                kelf_obs_13

#14-MER ELONGATION REACTIONS

DNA_1() -> DNA_15()                kelf_obs_14
```

```
DNA_2() -> DNA_16()                 kelf_obs_14

#15-MER ELONGATION REACTIONS

DNA_1() -> DNA_16()                 kelf_obs_15

end reaction rules

begin observables

Molecules TotalRad51
 ↪  DNA_1(),DNA_2(),DNA_3(),DNA_4(),DNA_5(),DNA_6(),DNA_7(),DNA_8()
,DNA_9(),DNA_10(),DNA_11(),DNA_12(),DNA_13(),DNA_14(),DNA_15(),DNA_16()

end observables

generate_network({overwrite=>1})
writeMfile({})
```

In the first section (`begin parameters`) all model parameters, including kinetic rate constants, are defined. In the second section (`begin molecule types`), all possible RAD51 polymer lengths on ssDNA and dsDNA are defined. In the third section (`begin seed species`), the initial condition for $S_0$ and $D_0$ is defined ($S_0(0) = D_0(0) =$ `DNA_start` $= 1$). In the fourth section (`begin reaction rules`), all allowed RAD51 polymerisation reactions are defined with their associated kinetic rate constants. The final section (`begin observables`) defines the desired model output (in this case $\sum_1^{16} iD_i$).

The resulting ODEs for the general ssDNA kinetic sub-model in compact form are as follows:

$$\frac{dS_0}{dt} = \sum_1^b \mathbf{k}_i^{\mathbf{d}} S_i - \mathbf{k_p} S_0 [\sum_1^b i(b+1-i)R_i] \quad (5)$$

$$\frac{dS_1}{dt} = -\mathbf{k}_1^{\mathbf{d}} S_1 + \mathbf{k_p} b R_1 S_0 - \mathbf{k_p} S_1 \sum_1^{b-1} R_i + \mathbf{k_s} S_2 \quad (6)$$

$$\frac{dS_2}{dt} = -\mathbf{k}_2^{\mathbf{d}} S_2 + \mathbf{k_p}[R_1 S_1 + 2(b-1)R_2 S_0] - \mathbf{k_p} S_2 \sum_1^{b-2} R_i + \mathbf{k_s}(S_3 - S_2) \quad (7)$$

for $3 \leq n \leq 15$:

$$\frac{dS_n}{dt} = -\mathbf{k}_n^{\mathbf{d}} S_n + \mathbf{k_p}[\sum_1^{n-1} R_i S_{n-i} + n(b+1-n)R_n S_0] - \mathbf{k_p} S_n \sum_1^{b-n} R_i + \mathbf{k_s}(S_{n+1} - S_n) \quad (8)$$

$$\vdots$$

$$\frac{dS_{16}}{dt} = -\mathbf{k}_{16}^{\mathbf{d}} S_{16} + \mathbf{k_p}[\sum_1^{15} R_i S_{16-i} + 16(b-15)R_{16} S_0] - \mathbf{k_s} S_{16} \quad (9)$$

With initial conditions $S_0(0) = 1, S_{1,\dots,16}(0) = 0$.

Similarly, the ODEs for the general dsDNA kinetic sub-model in compact form are as follows:

$$\frac{dD_0}{dt} = \sum_1^b \mathbf{k}_i^{\mathbf{d}} D_i - \mathbf{k_p} D_0 [\sum_1^b i(b+1-i)R_i] \tag{10}$$

$$\frac{dD_1}{dt} = -\mathbf{k}_1^{\mathbf{d}} D_1 + \mathbf{k_p} b R_1 D_0 - \mathbf{k_p} D_1 \sum_1^{b-1} R_i + \mathbf{k_s} D_2 \tag{11}$$

$$\frac{dD_2}{dt} = -\mathbf{k}_2^{\mathbf{d}} D_2 + \mathbf{k_p}[R_1 D_1 + 2(b-1)R_2 D_0] - \mathbf{k_p} D_2 \sum_1^{b-2} R_i + \mathbf{k_s}(D_3 - D_2) \tag{12}$$

for $3 \leq n \leq 15$:

$$\frac{dD_n}{dt} = -\mathbf{k}_n^{\mathbf{d}} D_n + \mathbf{k_p}[\sum_1^{n-1} R_i D_{n-i} + n(b+1-n)R_n D_0] - \mathbf{k_p} D_n \sum_1^{b-n} R_i + \mathbf{k_s}(D_{n+1} - D_n) \tag{13}$$

$$\vdots$$

$$\frac{dD_{16}}{dt} = -\mathbf{k}_{16}^{\mathbf{d}} D_{16} + \mathbf{k_p}[\sum_1^{15} R_i D_{16-i} + 16(b-15)R_{16} D_0] - \mathbf{k_s} D_{16} \tag{14}$$

With initial conditions $D_0(0) = 1, D_{1,\ldots,16}(0) = 0$.

$S_n$ and $D_n$ ($n = 1, ..., 16$) represent the time-dependent concentration of ssDNA-bound and dsDNA-bound RAD51 $n$-mers on the SPR chip, respectively. $S_0$ and $D_0$ represent the concentration of unbound ssDNA oligos and dsDNA oligos on the SPR chip, respectively. The adsorption of a RAD51 $i$-mer onto a DNA molecule ($i \leq b$) can occur on any $b+1-i$ sequence of $i$ DNA binding sites. In addition, the adsorption of a RAD51 $i$-mer can initiate via any of its $i$ protomers first binding to its corresponding DNA binding site with any of the $b+1-i$ sequences, and therefore the terms $\mathbf{k_p} S_0 R_i$ (5 - 9) and $\mathbf{k_p} D_0 R_i$ (10 - 14) are multiplied by $i(b+1-i)$. The $R_n$ values are constants that represent the concentration of RAD51 $n$-mers in solution and are derived from the equilibrium sub-model (1 - 4). At the end of the RAD51 SPR injection, all $R_n$ values are set to $0$. $\mathbf{k}_n^{\mathbf{d}}$ $(1 \leq n \leq 16)$, $\mathbf{k_p}$, and $\mathbf{k_s}$ are the kinetic rate constants describing the dynamics of the ODE models. $\mathbf{k}_n^{\mathbf{d}}$ represents the adsorption reverse rate constant for each $n$-mer on a DNA molecule with $b$ DNA binding sites (i.e. the one-step dissociation of RAD51 $n$-mers from DNA), $\mathbf{k_p}$ is the polymerisation rate constant, and describes the rate of adsorption and elongation of each $n$-mer on DNA, and $\mathbf{k_s}$ is the rate of RAD51 single protomer dissociation from DNA. The values of these rate constants are obtained by fitting the ODE model outputs ($\sum_1^b iS_i$ and $\sum_1^b iD_i$) to their respective data sets (Table 1) either via ABC-SMC or least-squares fitting. The initial conditions $S_0(0)$ and $D_0(0)$ were set to $1$ to match the SPR data normalisation, such that $(\max \sum_1^{16} iS_i) = (\max \sum_1^{16} iD_i) = 16$ (see Methods section in main text).

#### 1.4 Model Selection

The general ssDNA and dsDNA and kinetic sub-models (equations 5 - 9, equations 10 - 14) were used as templates to generate a set of ssDNA and dsDNA kinetic sub-models. In particular, we started from the simplest possible ssDNA and dsDNA kinetic sub-models that describe RAD51 polymerisation on DNA via 3 fit parameters: $\mathbf{k_p}$, $\mathbf{K_D}$, and $\mathbf{k_s}$ acting as both the single RAD51 protomer dissociation rate constant and the reverse adsorption rate constant for all RAD51 $n$-mers (KSM A: Table 2). We then generated further kinetic sub-models by either 1) sequentially increased the number of different adsorption reverse rate constants, or 2) varying the reactions regulated by specific adsorption reverse rate constants (KSMs B-I: Table 2).

Table 2: The set of ssDNA and dsDNA kinetic sub-models derived from the general ssDNA kinetic sub-model and the dsDNA kinetic sub-model. Rows describe the simplifications applied to the general ssDNA and dsDNA kinetic sub-models (equations 5 - 9, equations 10 - 14) to obtain sub-models A-I, and the total number of fit parameters for each sub-model. Total fit parameter counts include the $\mathbf{K_D}$ from the equilibrium sub-model. NA: not applicable. KSM: kinetic sub-model. ESM: equilibrium sub-model.

| KSM | ssDNA | Fit Parameters | dsDNA | Fit Parameters |
|---|---|---|---|---|
| A | $\mathbf{k}^{\mathbf{d}}_{1...16} = \mathbf{k_s}$ | 3: $\mathbf{k_s}$, $\mathbf{k_p}$, $\mathbf{K_D}$ | $\mathbf{k}^{\mathbf{d}}_{1...16} = \mathbf{k_s}$ | 3: $\mathbf{k_s}$, $\mathbf{k_p}$, $\mathbf{K_D}$ |
| B | $\mathbf{k}^{\mathbf{d}}_{1...3} = \mathbf{k_u}$ $\mathbf{k}^{\mathbf{d}}_{4...16} = \mathbf{k_s}$ | 4: $\mathbf{k_s}$, $\mathbf{k_u}$, $\mathbf{k_p}$, $\mathbf{K_D}$ | $\mathbf{k}^{\mathbf{d}}_{1} = \mathbf{k_u}$ $\mathbf{k}^{\mathbf{d}}_{2...16} = \mathbf{k_s}$ | 4: $\mathbf{k_s}$, $\mathbf{k_u}$, $\mathbf{k_p}$, $\mathbf{K_D}$ |
| C | $\mathbf{k}^{\mathbf{d}}_{1...3} = \mathbf{k_u}$ $\mathbf{k}^{\mathbf{d}}_{4,5} = \mathbf{k_s}$ $\mathbf{k}^{\mathbf{d}}_{6...16} = 0$ | 4: $\mathbf{k_s}$, $\mathbf{k_u}$, $\mathbf{k_p}$, $\mathbf{K_D}$ | $\mathbf{k}^{\mathbf{d}}_{1} = \mathbf{k_u}$ $\mathbf{k}^{\mathbf{d}}_{2} = \mathbf{k_s}$ $\mathbf{k}^{\mathbf{d}}_{3...16} = 0$ | 4: $\mathbf{k_s}$, $\mathbf{k_u}$, $\mathbf{k_p}$, $\mathbf{K_D}$ |
| D | $\mathbf{k}^{\mathbf{d}}_{1...3} = \mathbf{k_u}$ $\mathbf{k}^{\mathbf{d}}_{4} = \mathbf{k_q}$ $\mathbf{k}^{\mathbf{d}}_{5...16} = \mathbf{k_s}$ | 5: $\mathbf{k_s}$, $\mathbf{k_u}$, $\mathbf{k_q}$, $\mathbf{k_p}$, $\mathbf{K_D}$ | $\mathbf{k}^{\mathbf{d}}_{1} = \mathbf{k_u}$ $\mathbf{k}^{\mathbf{d}}_{2...16} = 0$ | 4: $\mathbf{k_s}$, $\mathbf{k_u}$, $\mathbf{k_p}$, $\mathbf{K_D}$ |
| E | $\mathbf{k}^{\mathbf{d}}_{1...3} = \mathbf{k_u}$ $\mathbf{k}^{\mathbf{d}}_{4} = \mathbf{k_q}$ $\mathbf{k}^{\mathbf{d}}_{5} = \mathbf{k_s}$ $\mathbf{k}^{\mathbf{d}}_{6...16} = 0$ | 5: $\mathbf{k_s}$, $\mathbf{k_u}$, $\mathbf{k_q}$, $\mathbf{k_p}$, $\mathbf{K_D}$ | $\mathbf{k}^{\mathbf{d}}_{1} = \mathbf{k_u}$ $\mathbf{k}^{\mathbf{d}}_{2} = \mathbf{k_q}$ $\mathbf{k}^{\mathbf{d}}_{3...16} = \mathbf{k_s}$ | 5: $\mathbf{k_s}$, $\mathbf{k_u}$, $\mathbf{k_q}$, $\mathbf{k_p}$, $\mathbf{K_D}$ |
| F | $\mathbf{k}^{\mathbf{d}}_{1...3} = \mathbf{k_u}$ $\mathbf{k}^{\mathbf{d}}_{4} = \mathbf{k_q}$ $\mathbf{k}^{\mathbf{d}}_{5...16} = 0$ | 5: $\mathbf{k_s}$, $\mathbf{k_u}$, $\mathbf{k_q}$, $\mathbf{k_p}$, $\mathbf{K_D}$ | $\mathbf{k}^{\mathbf{d}}_{1} = \mathbf{k_u}$ $\mathbf{k}^{\mathbf{d}}_{2} = \mathbf{k_q}$ $\mathbf{k}^{\mathbf{d}}_{3} = \mathbf{k_s}$ $\mathbf{k}^{\mathbf{d}}_{4...16} = 0$ | 5: $\mathbf{k_s}$, $\mathbf{k_u}$, $\mathbf{k_q}$, $\mathbf{k_p}$, $\mathbf{K_D}$ |
| G | $\mathbf{k}^{\mathbf{d}}_{1...3} = \mathbf{k_u}$ $\mathbf{k}^{\mathbf{d}}_{4} = \mathbf{k_q}$ $\mathbf{k}^{\mathbf{d}}_{5...16} = \mathbf{k_{r1}}$ | 6: $\mathbf{k_s}$, $\mathbf{k_u}$, $\mathbf{k_q}$, $\mathbf{k_{r1}}$, $\mathbf{k_p}$, $\mathbf{K_D}$ | $\mathbf{k}^{\mathbf{d}}_{1} = \mathbf{k_u}$ $\mathbf{k}^{\mathbf{d}}_{2} = \mathbf{k_q}$ $\mathbf{k}^{\mathbf{d}}_{3...16} = \mathbf{k_{r1}}$ | 6: $\mathbf{k_s}$, $\mathbf{k_u}$, $\mathbf{k_q}$, $\mathbf{k_{r1}}$, $\mathbf{k_p}$, $\mathbf{K_D}$ |
| H | $\mathbf{k}^{\mathbf{d}}_{1...3} = \mathbf{k_u}$ $\mathbf{k}^{\mathbf{d}}_{4} = \mathbf{k_q}$ $\mathbf{k}^{\mathbf{d}}_{5} = \mathbf{k_{r1}}$ $\mathbf{k}^{\mathbf{d}}_{6...16} = 0$ | 6: $\mathbf{k_s}$, $\mathbf{k_u}$, $\mathbf{k_q}$, $\mathbf{k_{r1}}$, $\mathbf{k_p}$, $\mathbf{K_D}$ | $\mathbf{k}^{\mathbf{d}}_{1} = \mathbf{k_u}$ $\mathbf{k}^{\mathbf{d}}_{2} = \mathbf{k_q}$ $\mathbf{k}^{\mathbf{d}}_{3} = \mathbf{k_{r1}}$ $\mathbf{k}^{\mathbf{d}}_{4...16} = 0$ | 6: $\mathbf{k_s}$, $\mathbf{k_u}$, $\mathbf{k_q}$, $\mathbf{k_{r1}}$, $\mathbf{k_p}$, $\mathbf{K_D}$ |
| I | $\mathbf{k}^{\mathbf{d}}_{1...3} = \mathbf{k_u}$ $\mathbf{k}^{\mathbf{d}}_{4} = \mathbf{k_q}$ $\mathbf{k}^{\mathbf{d}}_{5} = \mathbf{k_{r1}}$ $\mathbf{k}^{\mathbf{d}}_{6...16} = \mathbf{k_{r2}}$ | 7: $\mathbf{k_s}$, $\mathbf{k_u}$, $\mathbf{k_q}$, $\mathbf{k_{r1}}$, $\mathbf{k_{r2}}$, $\mathbf{k_p}$, $\mathbf{K_D}$ | NA | NA |

To fit the model to the data, we used an implementation of ABC-SMC [7] with a distance function defined by the sum-of-squared-residuals (SSR). To identify a point estimate of the parameters, we used the modes of the distributions. Each ssDNA kinetic sub-model was individually fit to the WT RAD51 ssDNA dN-X data (ssDNA dN-8, dN-14, dN-17, dN-50), and each dsDNA kinetic sub-model was individually fit to the WT RAD51 dsDNA dN-Xp data (dsDNA dN-5p, dN-8p, dN-11p, dN-50p).

The Akaike Information Criterion (AIC) is considered a standard tool used for model selection. In the case of sum-of-squares fitting, the AIC for each model can be calculated using the following equation [1]:

$$AIC = x \log\left(\frac{\text{SSR}}{x}\right) + 2p \tag{15}$$

Where $x$ represents the number of data points, SSR represents the sum-of-squared residuals, and $p$ represents the number of parameters in the model. Specifically, the AIC score is a trade-off between fit quality (i.e. the SSR value) and simplicity (i.e. the number of parameters $p$). Models are ranked from lowest AIC to highest AIC, and the model with the lowest AIC score best describes the data with the lowest number of parameters. In our scenario, each SPR curve consisted of 12512 data points, and each model was simultaneously fit to four SPR curves, meaning that the total $x$ was $x = 4(12512) = 50048$ data points. In this situation, the SSR and the number of data points strongly contribute to the AIC score, while the penalty factor $2p$ $(3 \leq p \leq 7)$ has a negligible impact on the AIC score (i.e. $AIC \approx x \log\left(\frac{\text{SSR}}{x}\right)$). As a result, the AIC is not sensitive to the number of parameters and given that the number of data points did not vary, we simply used the SSRs directly to identify the best-fit ssDNA sub-model (F - Table 2) (Fig. 2A in main text) and dsDNA sub-model (D - Table 2)

(Fig. 2B in main text) (Fig. 1).

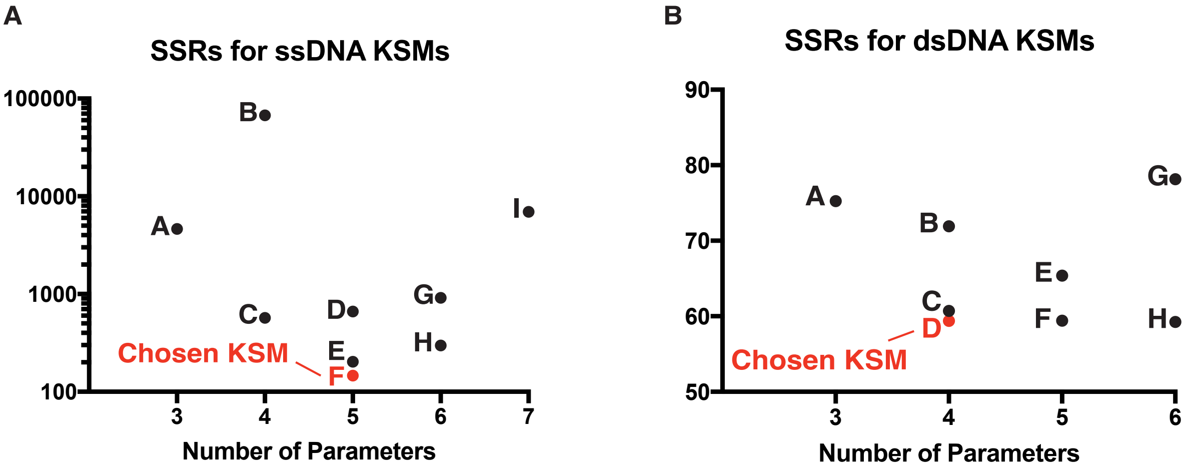


Figure 1: Selection of ssDNA and dsDNA kinetic sub-models. **A**. Sum-of-squared residuals (SSRs) for the ssDNA kinetic sub-models. Each kinetic sub-model together with the equilibrium sub-model was fit to one repeat of the ssDNA dN-X datasets via ABC-SMC. The chosen kinetic sub-model is indicated in red. **B**. SSRs for the dsDNA kinetic sub-models. Each kinetic sub-model together with the equilibrium sub-model was fit to one repeat of the dsDNA dN-Xp datasets via ABC-SMC. The chosen kinetic sub-model is indicated in red. A-B. KSM: kinetic sub-model.

The ssDNA kinetic sub-model F (Fig. 2A in main text) and dsDNA kinetic sub-model D (Fig. 2B in main text) were then simultaneously fit to the ssDNA dN-X data and the dsDNA dN-Xp data to obtain best fit parameter values (Fig. 2A-B in main text). Although the $\mathbf{k_u}$ fit for ssDNA dN-X was undetermined (Fig. 2A in main text), the probability density of $\mathbf{k_u}$ derived from the ABC-SMC fit (Fig. S2 in main text) nonetheless suggests dN-X $\mathbf{k_u} \gg$ dN-X $\mathbf{k_s}$. For this reason, the ssDNA kinetic sub-model F was further simplified to remove the $\mathbf{k_s} S_2$ and $\mathbf{k_s} S_3$ terms from equations 6 - 8.

ABC-SMC simulations were also carried out in MATLAB 2016b to simultaneously fit the F86E RAD51 ssDNA dT-50, dN-50, dA-50 and the dsDNA dN-50p SPR data using the ssDNA and dsDNA kinetic sub-model A (Fig. 4D in main text).

### References

[1] K P Burnham and D R Anderson. *Model Selection and Multimodel Inference - A Practical Information-Theoretic Approach*. Springer, 2002.
